# Systemic extracellular acidification is a hallmark of aging

**DOI:** 10.1101/2024.09.24.614672

**Authors:** Eliano dos Santos, Yining Xie, Enyuan Cao, Andrea Foley, Max E. Taylor, Ivan Andrew, George Young, Natalie L. Trevaskis, Helena M. Cochemé

## Abstract

Understanding the critical pathophysiological processes that promote age-related disease is needed to uncover effective targets for preventive medicine. Here, we investigate how extracellular pH changes with age and its impact on longevity, using fly and mouse models. We find that extracellular acidification occurs in flies during aging and correlates to mortality rate. With age, flies also become more susceptible to die from acidotic stress, which can be prevented by alkalotic treatment. Acidification is caused by insufficient acid elimination, linked to downregulation of genes in the fly excretory tract that control pH and ATP production, essential for active secretion initiation. In mice, we show that lymph-drained interstitial fluids acidify with age. Expression of genes, whose pathogenic loss-of-function variants cause tubular acidosis in humans, is decreased in the kidneys of aging mice. Overall, this study sheds light on dysregulated systemic acid-base balance as a conserved pathophysiological mechanism of aging.

## Introduction

The incidence of chronic diseases and geriatric syndromes as populations age is a growing public health challenge, with severe economic and societal consequences^1,2^. Aging is the main risk factor for human frailty and related chronic diseases^3,4^, yet our understanding of the pathophysiological processes promoting this risk is incomplete.

Categorising age-related processes as hallmarks of aging prompted the discovery of novel mechanisms and therapeutic targets for preventive medicine in old age^5^, but the fundamental causes of aging remain contentious^6^. In the hallmarks classification, primary hallmarks that reflect progressive accumulation of damage with age are interpreted as causal because they occur at the lowest level of biological organization from all considered: molecules^5,7^. Taking this reductionist view further, the behavior of subatomic particles can be proposed as an upstream mechanism.

Both protons and electrons alter the structure and function of molecules. Electrons have been thoroughly studied in aging, originally as the focus of the free radical theory of aging^8^, and more recently as an integral part of intracellular redox signalling^9^. In comparison, protons have received little attention by geroscience. Despite their very low concentration in bodily fluids, hereafter referred to as pH (i.e. −log[H^+^]), protons have very high charge density, and determine the structure and configuration of macromolecules in their proximity^10^. By influencing the structure and activity of proteins, including enzymes, protons effectively control all biological processes.

Given its key role in biology, pH is tightly regulated across all bodily fluid compartments. Here, we focus on the extracellular pH as a representation of systemic acid-base balance. Human blood pH typically ranges from 7.35 to 7.45, and small fluctuations can significantly impact physiology^11^. As an essential and redundantly regulated property, blood pH would be expected to remain constant throughout life, yet a sub-clinical metabolic acidification occurs in humans as they age^12^. These seemingly small changes in blood pH likely reflect a more profound systemic acidification, particularly in interstitial fluids with lower immediate compensatory capacity. For example, localised acidity within an organ can escape prompt systemic compensation, as interstitial fluids from both affected and unaffected regions are jointly drained and buffered before joining systemic circulation.

Interstitial fluids are challenging to collect, so their acid-base physiology at the tissue level is unexplored, particularly with age. Here, we use *Drosophila melanogaster* (henceforth referred to as *Drosophila* or fly) as the primary model to study the impact of acid-base balance on longevity. Besides having a comparatively short lifespan to mammals, strong evolutionary conservation of metabolic pathways implicated in aging, and powerful genetic tools^13,14^, the fly is particularly useful to study systemic extracellular fluid pH with age. *Drosophila* has an open circulatory system^15^, with a single extracellular fluid compartment, the hemolymph, bathing all organs (Fig. 1a), making it a simpler model to manipulate and directly correlate extracellular pH changes with whole-organism readouts. Flies also have complex organs which are orthologous to their mammalian counterparts. These include an excretory tract that comprises Malpighian tubules, functionally comparable to renal tubules, which regulate acid-base balance^16^.

**Fig 1.**
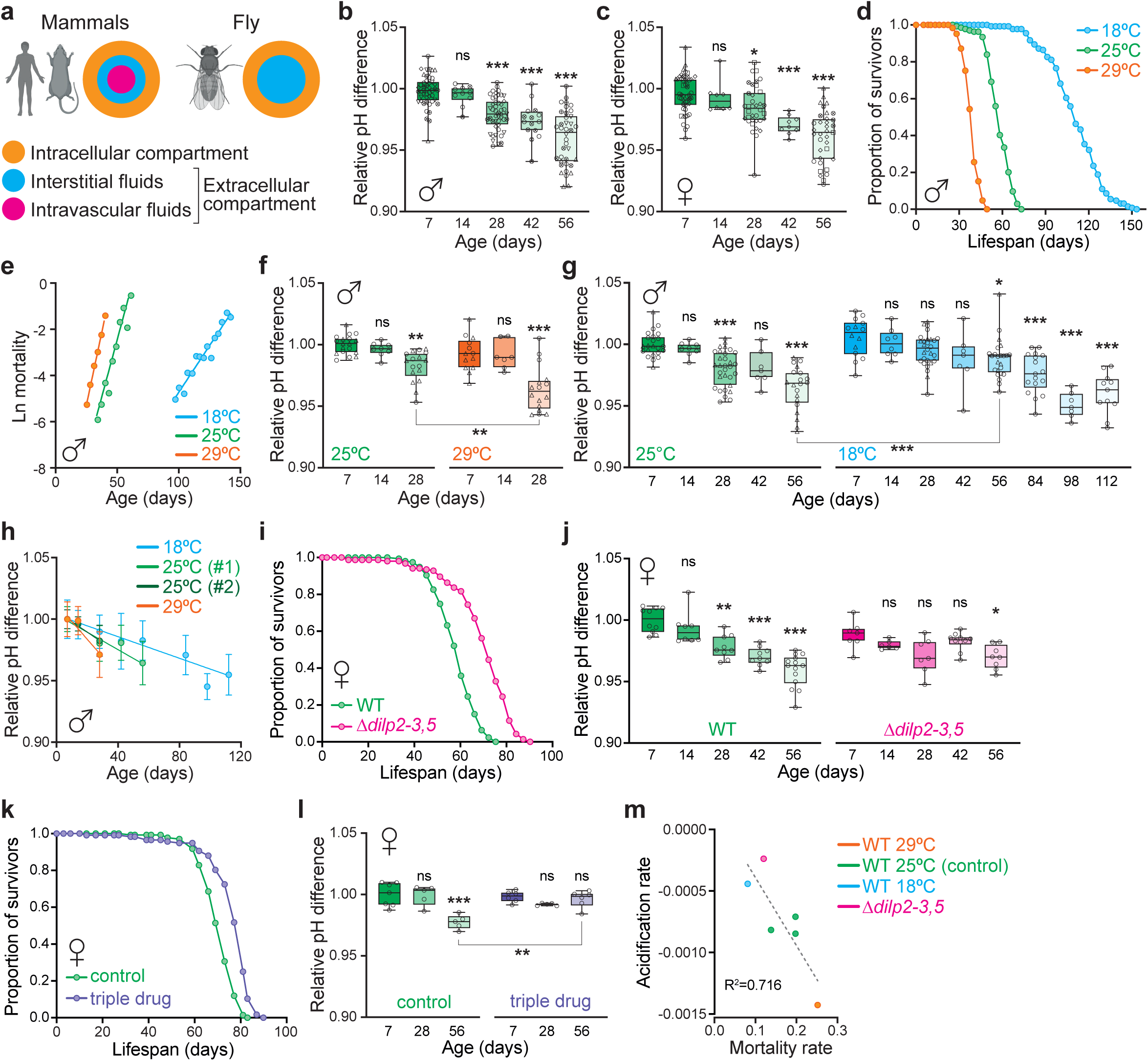
Extracellular acidification of adult *Drosophila* correlates with mortality rate. **a,** The extracellular compartment in mammals is divided into intra- and extravascular (interstitial) fluids, while *Drosophila* has a single extracellular fluid, the hemolymph. **b-c,** Hemolymph pH differences with age in WT (*w^Dah^*) flies, raised under standard 25°C conditions, relative to the young (7-day old, d7) control. Data for males (**b**) and females (**c**) (n=8-48 total replicates per condition, pooled from 5 and 7 independent experiments, respectively). **d-e,** Survival of male WT (*w^Dah^*) flies raised at 18, 25 and 29°C. Lifespan curves (**d**, n=127-136 flies per condition). Mortality rates (**e**, slopes: 25°C v. 29°C, p=0.16; 25°C v. 18°C, p<0.001). **f-h,** Hemolymph pH differences with age in male WT (*w^Dah^*) flies raised at different temperatures, relative to the d7 25°C control. 25°C and 29°C (**f**, n=7-18 total replicates per condition, pooled from 2 independent experiments). 25°C and 18°C (**g**, n=7-31 total replicates per condition, pooled from 3 independent experiments). Rates of hemolymph acidification, relative to the d7 controls for each temperature (**h**). Data are means ±SD (slopes: 25°C v. 29°C, p=0.06; 25°C v. 18°C, p<0.01). **i-j,** Analysis of the IIS pathway mutant *Δdilp2-3,5* against female WT (*w^Dah^*) controls. Lifespan curves (**i**, n=154-157 flies per condition). Hemolymph pH differences with age, relative to the d7 WT control (**j**, n=6-14 total replicates per genotype). **k-l,** Response of female WT (*w^Dah^*) flies treated with a triple drug combination of 50 μM rapamycin, 15.6 μM trametinib and 1 mM lithium. Lifespan curves (**k**, n=117-135 flies per condition). Hemolymph pH differences with age (**l**, n=5-7 replicates per condition), relative to the untreated d7 control. **m,** Correlation between rates of mortality and hemolymph acidification across conditions (R^2^=0.716). Graphs in **b**, **c**, **f**, **g**, **j** and **l** are box-plots, with median and min-max error bars. Statistical significance was determined by one-way ANOVA (Tukey). Each pH data point is based on the hemolymph collected from a cohort of 15 male or 12 female flies, with different symbols corresponding to independent experiments. Unless otherwise indicated, statistical comparison is relative to the d7 control within each condition. Survival data in **d**, **i** and **k** were analyzed by Log-Rank test (see Supplementary Table 1 for full details of n numbers and p values). Slopes of mortality (**e**) and hemolymph acidification rates with age (**h**) were calculated by simple linear regression and compared by analysis of covariance. ns, p>0.05; *, p<0.05; **, p<0.01; ***, p<0.001.

Here, we establish a link between extracellular pH and fly mortality rate, and develop a model of metabolic acidosis in flies, which uncovers increased susceptibility to acidotic stress with age. We find no evidence for increased acid production in old flies, despite changes in metabolic rate with age. Instead, the extracellular compartment acidifies with age due to insufficient acid elimination, which relates to lower expression of most acid-base related genes expressed in the fly excretory tract. In agreement with these findings, lymph-drained interstitial fluid in mice also acidifies with age, and the expression of ortholog genes, with pathogenic variants known to cause metabolic acidosis in humans, decreases in the mouse kidney. Overall, this study highlights the importance of acid-base balance in aging, and implicates acidification as a fundamental, evolutionarily conserved pathophysiological mechanism that increases frailty and disease risk with age to ultimately limit lifespan.

## Results

### Extracellular acidification of adult *Drosophila* correlates with mortality rate

To test whether extracellular fluid acidification is a hallmark of aging, we measured the pH of extracted fly hemolymph with age using a cell-impermeable pH-sensitive pyranine dye (Extended Data Figs. 1a-c)^17^. Since the standard curve is made with buffer solutions rather than fly hemolymph, and the calculated pH is dependent on the composition and osmolarity of these buffer solutions, we focused on relative pH differences with age rather than absolute values (pH ∼7.1-7.2 for young flies under control conditions).

We aged wild-type (WT) flies at 25°C on standard sugar-yeast-agar (SYA) medium and collected their hemolymph at different timepoints ranging from young to old (days 7, 14, 28, 42 and 56 of adulthood). We found that fly hemolymph became significantly more acidic with age in both males and females (Figs. 1b,c). Age-related hemolymph acidification also occurred independently of sex in other fly strains (Extended Data Figs. 1d-g) and dietary conditions (Extended Data Fig. 1h).

To test whether extracellular acidification relates to mortality, we considered a range of interventions that are known to modulate lifespan in *Drosophila.* Firstly, we explored the effect of temperature. Flies are ectotherms, and altering the environmental temperature alters their mortality rate^18^. While control flies are typically maintained at 25°C, lifespan is significantly shortened when raised at 29°C, and conversely extended at 18°C (Figs. 1d,e). Since fluid pH is determined by temperature^19^, and temperature inversely correlates with both pH and lifespan, we tested whether the changes in mortality rate could be related to a direct life-long effect of temperature on hemolymph pH. We collected hemolymph from age-matched male WT flies at the different temperatures, and found that the rate of hemolymph acidification with age was faster at 29°C and slower at 18°C (Figs. 1f-h), reflecting the respective mortality rates. Notably, the hemolymph pH of the longer-lived 18°C flies eventually dropped as the mortality rate increased at later ages to levels comparable with old flies raised at 25°C (Figs. 1g,h), suggesting the effect of temperature on age-related acidification rate is indirect, i.e. through pathophysiological mechanisms.

Next, we investigated a genetic paradigm of longevity, using the long-lived *Δdilp2-3,5* mutant. *Drosophila* has 8 insulin-like peptides (dILPs) that are the ligands for the sole fly insulin receptor and activate the *Drosophila* insulin/IGF-like signalling (IIS) pathway. Genetic deletion of dILPs 2, 3 and 5, expressed in the fly brain median neurosecretory cells, significantly extended the lifespan of female flies (Fig. 1i and Extended Data Fig. 1i)^20^, consistent with the observation that downregulation of the IIS pathway promotes longevity in an evolutionary conserved manner across organisms, from yeast to mammals^21^. We collected hemolymph from Δ*dilp2-3,5* females and age-matched WT controls, and observed that systemic acidification occurred at a slower rate in the long-lived IIS mutant (Fig. 1j and Extended Data Fig. 1j). Finally, we tested a pharmacological mode of lifespan extension, treating flies throughout adulthood with a triple drug combination of rapamycin, trametinib and lithium, which acts by targeting nutrient-sensing pathways^22^. Similarly to the genetic IIS mutant, we found that long-lived triple drug-treated female flies exhibited less acidic hemolymph at older age (Figs. 1k,l and Extended Data Figs. 1k,l).

Collectively, these data show that systemic acidification with age occurs irrespective of sex, strain and diet, and that independent lifespan-extending interventions correlate with slower rates of hemolymph acidification (Fig. 1m). Therefore, altered pH homeostasis is a common pathophysiological process that ultimately occurs even in long-lived populations as their mortality rate increases. This suggests that systemic acidification is either a common consequence of aging, or a fundamental cause whose effects are ameliorated by different interventions.

### Acidotic stress resistance declines with age

To test whether physiological acidification can accelerate mortality, we fed flies ethylene glycol (EG) to induce metabolic acidosis. EG is metabolized into glycolaldehyde by alcohol dehydrogenase before being converted into glycolic acid by aldehyde dehydrogenases^23^, and causes acute metabolic acidosis in humans^11,24^. As an acid precursor, EG did not alter the pH of our standard fly media (pH ∼4.5). Young WT flies were exposed to EG from day 2 of adulthood at a range of concentrations (0.05 to 5%), and hemolymph was collected after 7 days of treatment. Hemolymph pH decreased upon EG treatment in a dose-dependent manner, independently of sex or diet (Fig. 2a and Extended Data Fig. 2a). The highest dose (5% EG) was lethal before the hemolymph collection timepoint (Figs. 2b,c). 2.5% EG caused a drop in hemolymph pH after 7 days comparable to the relative acidification with age while not being acutely lethal, and was therefore used for subsequent acidotic stress assays.

**Fig 2.**
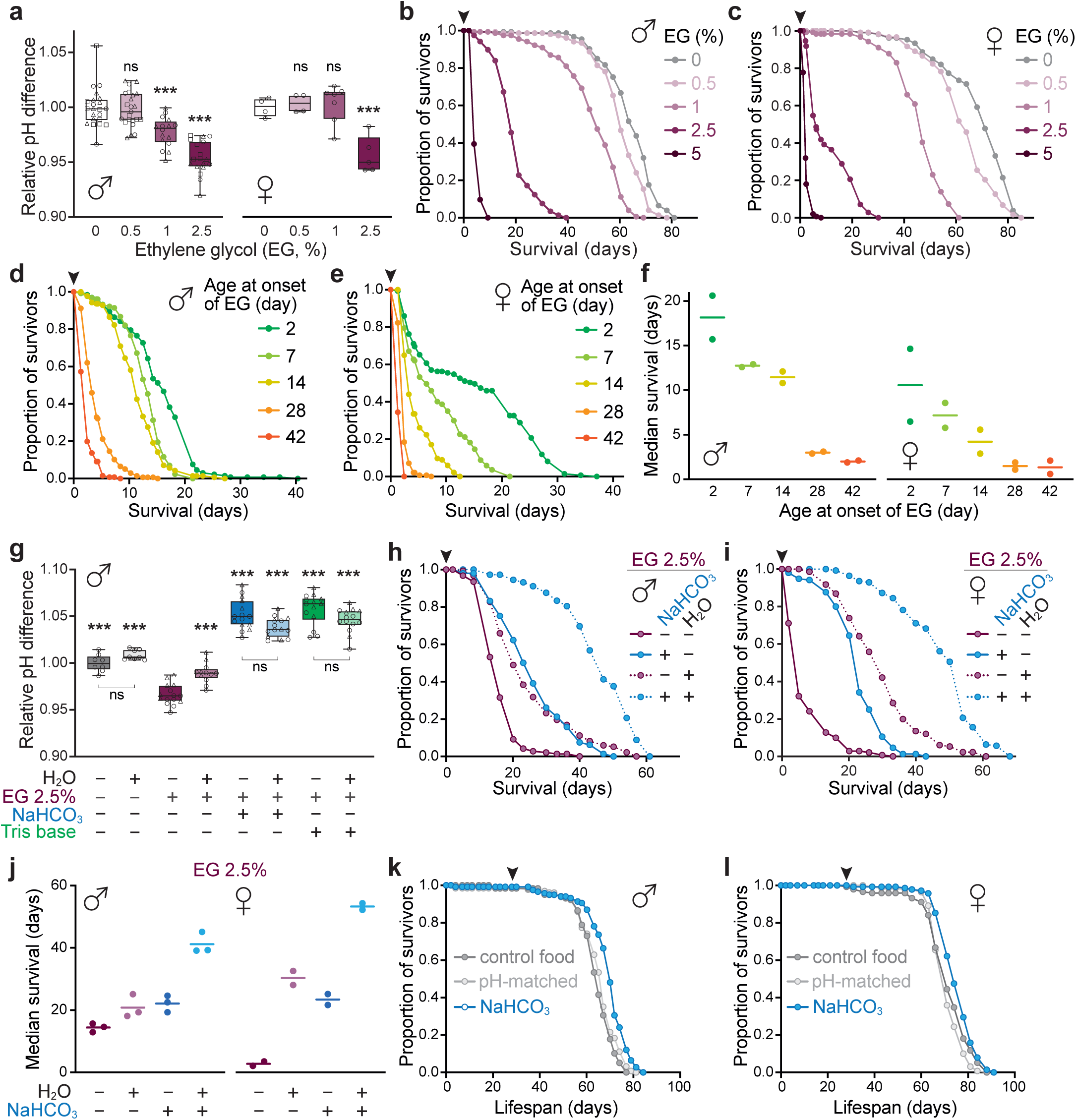
Susceptibility to acidotic stress increases with age. **a,** Hemolymph pH differences in male and female WT (*w^Dah^*) flies upon exposure to EG (0.5-2.5%) for 7 days, relative to the untreated control for each sex. Total n=18-23 replicates for males, pooled from 3 independent experiments, and n=7-8 for females. **b-c,** Survival curves of WT (*w^Dah^*) flies exposed to EG (0.5-5%) throughout their lifespan, from day 2 of adulthood (arrowhead). Males (**b**, n=148-158 per condition) and females (**c**, n=106-148 per condition). **d-f,** Response of WT (*w^Dah^*) flies to chronic 2.5% EG exposure initiated at different ages (from day 2, 7, 14, 28 and 42 of adulthood). Representative survival curves of males (**d**, n=129-141) and females (**e**, n=134-145). Arrowheads indicate the start of treatment. Summary of median survival data for 2 independent experiments (**f**). **g,** Hemolymph relative pH differences of WT (*w^Dah^*) males exposed to 2.5% EG and treated with 100 mM NaHCO_3_ or Tris base for 7 days, with and without an extra source of water, compared to control flies (n=8-15 replicates per condition, pooled from 2 independent experiments). Statistical comparison is against the 2.5% EG condition. **h-j,** Effect of 100 mM NaHCO_3_ and water supplementation on the survival of WT (*w^Dah^*) flies exposed to 2.5% EG throughout their lifespan, from day 2 of adulthood (arrowhead). Representative survival curves of males (**h**, n=151-158 per condition) and females (**i**, n=142-155 per condition). Summary of median survival data for 2 independent experiments (**j**). **k-l,** Survival curves of WT (*w^Dah^*) flies treated with 100 mM NaHCO_3_ continuously from middle-age (day 28 of adulthood, indicated by arrowhead). Males (**k**, n=124-133 per condition) and females (**l**, n=142-147 per condition). Graphs in **a** and **g** are box-plots, with median and min-max error bars. Statistical significance was determined by one-way ANOVA (Tukey). Each pH data point is based on the hemolymph collected from a cohort of 15 male or 12 female flies, with different symbols corresponding to independent experiments. Survival data in **b**-**e**, **h**, **i**, **k** and **l** were analyzed by Log-Rank test (see Supplementary Table 1 for full details of n numbers and p values). ns, p>0.05; ***, p<0.001.

At lower doses, EG has been used to induce tubule lithiasis in animal models^23,25^. Indeed, patients often have oxalate crystals in their urine after EG poisoning^24^. Glycolic acid can be metabolized into oxalic acid, which almost fully dissociates at physiological pH into oxalate and H^+^. Oxalate is filtered by the kidney, where it can bind to calcium and crystalize to induce lithiasis. EG is reported to cause deposits of calcium oxalate crystals in the fly Malpighian tubules in a dose-dependent manner up to 1%^25^. Acute tubular damage caused by calcium oxalate crystals could therefore contribute to the metabolic acidosis associated with EG. Unexpectedly, we did not observe crystal formation in the tubules of male flies after 7 or 14 days of 2.5% EG feeding, when hemolymph pH is already significantly decreased (score of 0 based on n=20 dissected *w^Dah^* WT male flies per condition)^26^.

To further explore excretory dysfunction as a potential contributor to EG-induced acidosis independently of crystal deposition, we analyzed fly excreta by adding a pH-sensitive dye to the food (Extended Data Fig. 2b). Fly excreta consist of urine, produced by the tubules and drained into the hindgut, combined with digestive waste. By measuring excreta pH, we are able to assess the total fixed acid excretion over a period of time. In response to 2.5% EG treatment, young flies produced more deposits, which tended to be bigger (Extended Data Figs. 2c,d), and importantly had a lower pH (Extended Data Fig. 2e). Overall, this suggests that the fly excretory tract is still efficiently excreting acid upon acidotic stress, but the rate of excretion is insufficient to maintain acid-base balance.

Importantly, we found that acidosis induced by EG caused a dose-dependent shortening of fly lifespan, independently of sex or strain (Figs. 2b,c and Extended Data Figs. 2f,g). To test the susceptibility to acidotic stress with age, we treated flies with 2.5% EG at different ages, and observed that older flies were increasingly more sensitive to acidotic stress (Figs. 2d-f). This enhanced sensitivity occurred despite flies eating less as they age^27^, suggesting that even lower intake of the same EG dose is sufficient to cause early acidosis-related mortality at older ages.

### Alkalotic treatment counteracts acidotic stress and extends fly lifespan

Since chronic acidotic stress decreased fly survival, we next aimed to rescue survival under acidotic challenge by manipulating hemolymph pH pharmacologically. Alkalotic treatment can be an option to rapidly re-establish acid-base balance in cases of severe acidemia, or less severe acidemia associated with acute kidney injury in humans^28^. The standard treatment is sodium bicarbonate (NaHCO_3_), but tromethamine (Tris base) has also been used for that purpose^11^. These alkalotic compounds have different properties and mechanisms of action, so we tested both as independent buffers added to the fly food. EG acts as a strong osmole, which may cause dehydration upon excretion, so we also provided flies with an additional water source, as previously established^26^, and measured fly hemolymph pH after 7 days of treatment Under acidotic stress, both alkalotic treatments significantly increased hemolymph pH (Fig. 2g). Water did not have an additive effect to the alkalotic compounds, but alone rescued EG-induced acidification, suggesting that dehydration plays a role in limiting compensatory acid excretion. Importantly, both NaHCO_3_ and water alone extended the survival of male and female flies under acidotic stress (Figs. 2h-j). Notably, the combination of water with alkalotic treatments further rescued the lifespan of flies in both sexes, showing that acidification and dehydration are the main mechanisms shortening fly survival upon EG treatment. At a high concentration (100 mM), Tris base alone did not rescue fly survival under acidotic stress (Extended Data Fig. 2h), likely related to direct toxicity of this alkalotic agent considering the significant survival extension when combined with water. Indeed, 100 mM Tris base shortened fly lifespan under standard conditions (Extended Data Fig. S2i). However, a lower concentration (10 mM) extended lifespan of WT males (Extended Data Fig. 2j)^29^, without causing alkalosis in healthy, young flies (Extended Data Fig. 2k).

To explore the effects of alkalotic treatment on survival under standard conditions, we treated flies with NaHCO_3_ from middle-age (day 28 of adulthood), when fly hemolymph already exhibited acidification (Figs. 1b,c). We found that NaHCO_3_ treatment extended the lifespan of both male and females flies, independently of food pH (Figs. 2k,l). This effect was also observed with NaHCO_3_ supplementation throughout adulthood (Extended Data Figs. 2l,m). In the absence of osmotic stress, an extra source of water did not further extend lifespan (Extended Data Fig. 2l). Similar to Tris base, NaHCO_3_ treatment in young WT flies tended to only mildly alkalinize their hemolymph (Extended Data Fig. 2n). Therefore, healthy flies can maintain acid-base balance to prevent alkalosis in response to alkalotic agents under control conditions, whereas under acidotic stress the same dose of alkalotic treatment causes significant hemolymph alkalinization (Fig. 2g). Flies fed at the same rate or more when given NaHCO_3_ or pH-matched food (Extended Data Fig. 2o), excluding dietary restriction as a potential contributor to the survival effect.

Together, these results indicate that manipulating hemolymph pH can impact survival. However, alkalotic compounds are blunt tools that can have side-effects by disrupting pH balance towards alkalosis in healthy animals. Therefore, we focused on establishing the source of acid accumulation with age to gain insight into specific potential therapeutic strategies.

### Global metabolic production of acids does not increase with age

Acid can accumulate systemically, either because increased metabolic production overwhelms elimination, or due to retention by loss of excretory function (Extended Data Fig. 2p). To assess global volatile acid production with age in flies, we used high resolution *in vivo* respirometry to measure their total CO_2_ production and O_2_ consumption over ∼24 h (Fig. 3a). The ratio of produced CO_2_ to consumed O_2_ (also known as respiratory quotient, RQ) reflects changes in global substrate utilization and acid production. Theoretically, the RQ is predicted as ∼1 for carbohydrates, versus ∼0.7 for fats and ∼0.8-0.9 for proteins, depending on the fatty acid and amino acid composition, respectively^30^. Only metabolism of volatile organic acids (converted to CO_2_) can result in a RQ value above 1. As proof of principle, we treated young WT flies with EG for 7 days, and indeed observed an increased RQ (Fig. 3b), although being <1 a metabolic shift to higher consumption of carbohydrate may not be excluded. Interestingly, acidotic stress lowered metabolic rate (Extended Data Figs. 3a,b), which may be a compensatory mechanism to decrease acid production.

**Fig 3.**
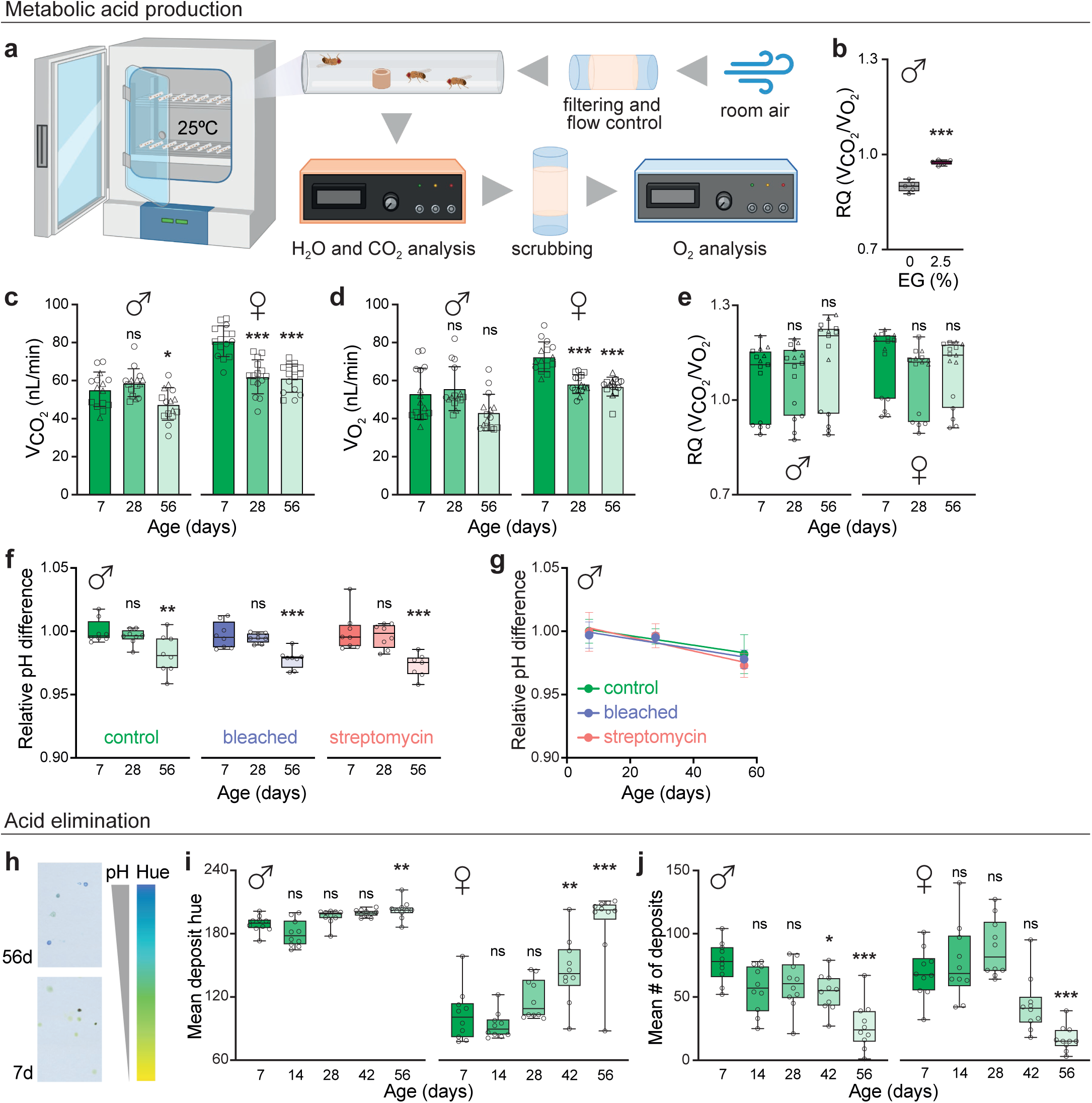
Global acid elimination declines with age in *Drosophila.* **a,** Scheme of the fly respirometry set-up. Room air is filtered of CO_2_, dehumidified and pumped into 2 valves that control air flow into the system. One valve is connected to the gas analyzers for direct reference air analysis. The other valve directs air through a multiplexer to chambers in a temperature-controlled incubator containing flies with access to food. In stop-flow cycles of 48 min, during which flies exchange gas within the chamber, the accumulated air progresses from each chamber into the CO_2_ and H_2_O analyzers and, after scrubbing, into the O_2_ analyzer. **b,** Mean respiratory quotient (RQ) per chamber of young (7d) WT (*w^Dah^*) males exposed to 0 or 2.5% EG for ∼24 h. Data are box-plots with median and min-max error bars (n=5 chambers per condition, with 3 flies), analyzed by unpaired two-tailed *t*-test. **c-e,** Respirometry analysis of mated male and female WT (*w^Dah^*) flies at different ages. Total n=15 replicates (chambers) per condition, pooled from 3 independent experiments with n=5 chambers per condition with 3 flies per chamber. Mean ±SD of CO_2_ production (**c**). Mean ±SD of O_2_ consumption (**d**). Mean RQ (box-plots with median and min-max error bars) (**e**). Statistical comparison is against the d7 within each condition, analyzed by one-way ANOVA (Tukey). **f-g,** Hemolymph pH differences with age in WT (*w^Dah^*) males upon microbiota deprivation, either through egg bleaching or streptomycin treatment. Data are box-plots with median and min-max error bars (n=7-8 replicates per condition), relative to the untreated 7d control with normal microbiota (**f**). Statistical comparison is against the d7 within each condition, analyzed by one-way ANOVA (Tukey). Rates of hemolymph acidification, relative to the 7d for each condition (**g**). Data are means ±SD (slopes are not significantly different, p=0.46). Each pH data point is based on the hemolymph collected from a cohort of 15 male flies. **h-j,** Analysis of excreta from male and female WT (*w^Dah^*) flies at different ages. Representative scans of excreta from young (7d) and old (56d) females upon feeding with the pH-sensitive dye, bromophenol blue (**h**). Lower hue (towards yellow) corresponds to more acidic deposits. Mean deposit hue (**i**), and mean number (**j**) of deposits. Data are box-plots with median and min-max error bars (n=10 plates, each with 5 flies). Statistical comparison is against the d7 for each sex, analyzed by one-way ANOVA (Tukey). ns, p>0.05; *, p<0.05; **, p<0.01; ***, p<0.001.

Next, we applied *in vivo* respirometry to assess global metabolic changes with age. In line with previous reports^31–33^, production of CO_2_ in male flies did change substantially with age (Fig. 3c and Extended Data Figs. 3c,i). However, CO_2_ alone is not a reliable measure of metabolic rate. For instance, lowering of metabolic rate could be masked by higher production and buffering of organic acids, and appear constant. Therefore, together with CO_2_, we comprehensively measured O_2_ consumption with age in flies of different genetic backgrounds (*w^Dah^* and *w^1118^*). Similarly to CO_2_, we found that O_2_ consumption did not significantly change in males, either during light or dark cycles (Fig. 3d and Extended Data Figs. 3d,j), which translated to no significant changes in RQ (Fig. 3c and Extended Data Figs. 3e,k). Conversely, female flies exhibited lower metabolic rate during aging, irrespective of mating status, and with different circadian patterns (Figs. 3c,d and Extended Data Figs. 3c-j), but no consistent increase in RQ across strains (Fig. 3e and Extended Data Figs. 3e,h,k). In all conditions, RQ was variable and tended to be >1 at all ages, indicating that volatile organic acids are produced throughout the fly lifespan, possibly by glycolytic tissues, such as muscle^34^. Importantly, RQ did not significantly increase with age, suggesting that the systemic acidification does not result from increased acid production, but rather loss of excretory capacity with age under continuous production of acids.

Besides volatile organic acids, altered absorption of acidogenic amino acids or gut microbial production of fixed acids could increase acid load with age. Indeed, the fly gut microbiota can change with age^35,36^. To explore the association between age-related hemolymph acidification and intestinal microbiota, flies were cleared of their microbiota, either by dietary supplementation with a broad-spectrum antibiotic, which can be administered chronically without impacting fly lifespan^37^, or through bleaching of the eggs. Both streptomycin treatment and bleaching were sufficient to effectively eliminate the fly gut microbiota (Extended Data Fig. 3l), as previously described^38^, and streptomycin treatment itself did not alter hemolymph pH (Extended Data Fig. 3m). Importantly, systemic age-related acidification occurred similarly under control and microbiota-deprived conditions (Figs. 3f,g), confirming that gut bacteria do not underlie this process.

### Acid elimination declines with age

Given the absence of global changes in acid production, we then tested whether hemolymph acidification with age occurred as a result of insufficient acid elimination. In flies, excretion starts at the Malpighian tubule, where tubule-secreted fluids are drained into the hindgut and join with digestive waste to ultimately be excreted as feces (Extended Data Fig. 2p). Unlike mammalian kidneys, fly tubules are not preceded by filtering units and their cells are joined by septate junctions that close the paracellular space^39^. Secretion is instead actively initiated by vacuolar-type ATPases (v-ATPases) at the apical membrane which pump protons into the tubule lumen, creating a H^+^ gradient that drives strong ion and water secretion^40,41^. Therefore, excretory function is intimately linked to ion channels that impact acid-base balance in flies.

To measure total acid excretion with aging, flies were fed a pH-sensitive dye at different ages throughout their lifespan and their excreta were analyzed over 24 h (Fig. 3h). Acidification of deposits is indicated by decreased hue, as the dye color changes towards yellow^42,43^. Since flies have a single combined waste product, analyzing the pH of their excreta captures total acid excretion over the 24 h period. As hemolymph acidifies with age (Figs. 1b,c), the excretory tract would be expected to compensate by promoting acid elimination in order to maintain homeostasis. Instead, excreta pH increased with age in both sexes (Fig. 3i), more prominently in females, while the number of deposits declined over time (Fig. 3j). This suggests that hemolymph acidification is a consequence of insufficient acid elimination, and is consistent with previous findings that Malpighian tubule secretion rates decline with age^44,45^.

To further analyze how excretory tract insufficiency occurs with age, we performed RNA-seq on the dissected excretory tracts (hindgut and tubules) of young, middle-aged and old male flies (days 7, 28 and 56 of adulthood), and observed global gene expression changes with age (Fig. 4a). By middle-age (28 days), the fly excretory tract already showed a significant downregulation of genes involved in ATP production by aerobic respiration (Fig. 4b). Mitochondria-derived ATP is essential for secretion to be initiated by v-ATPases. In the Malpighian tubules, mitochondria locate by the apical membrane, and inhibition of aerobic respiration-related ATP production prevents electrogenesis and transmembrane ion transport, inhibiting fluid secretion^40,41,46^. Surprisingly, higher cytosolic levels of ATP were recently described in the principal tubule cells of middle-aged flies compared to young, despite signs of mitochondrial dysfunction^45^. We found no clear compensation by upregulation of genes related to alternative ATP production routes in the excretory tract of either middle-aged or old flies (Extended Data Figs. 4a-c). Glucose uptake was also shown to decline in the tubular principal cells of middle-aged flies^45^. In yeast and mammalian renal epithelial cells, v-ATPase assembly is dependent on intracellular glucose concentrations^47^. Given v-ATPase is a major consumer of ATP for secretion initiation, primary v-ATPase loss-of-function associated with lower glucose uptake may explain higher levels of cytosolic ATP, despite lower production by mitochondria at the apical membrane.

**Fig 4.**
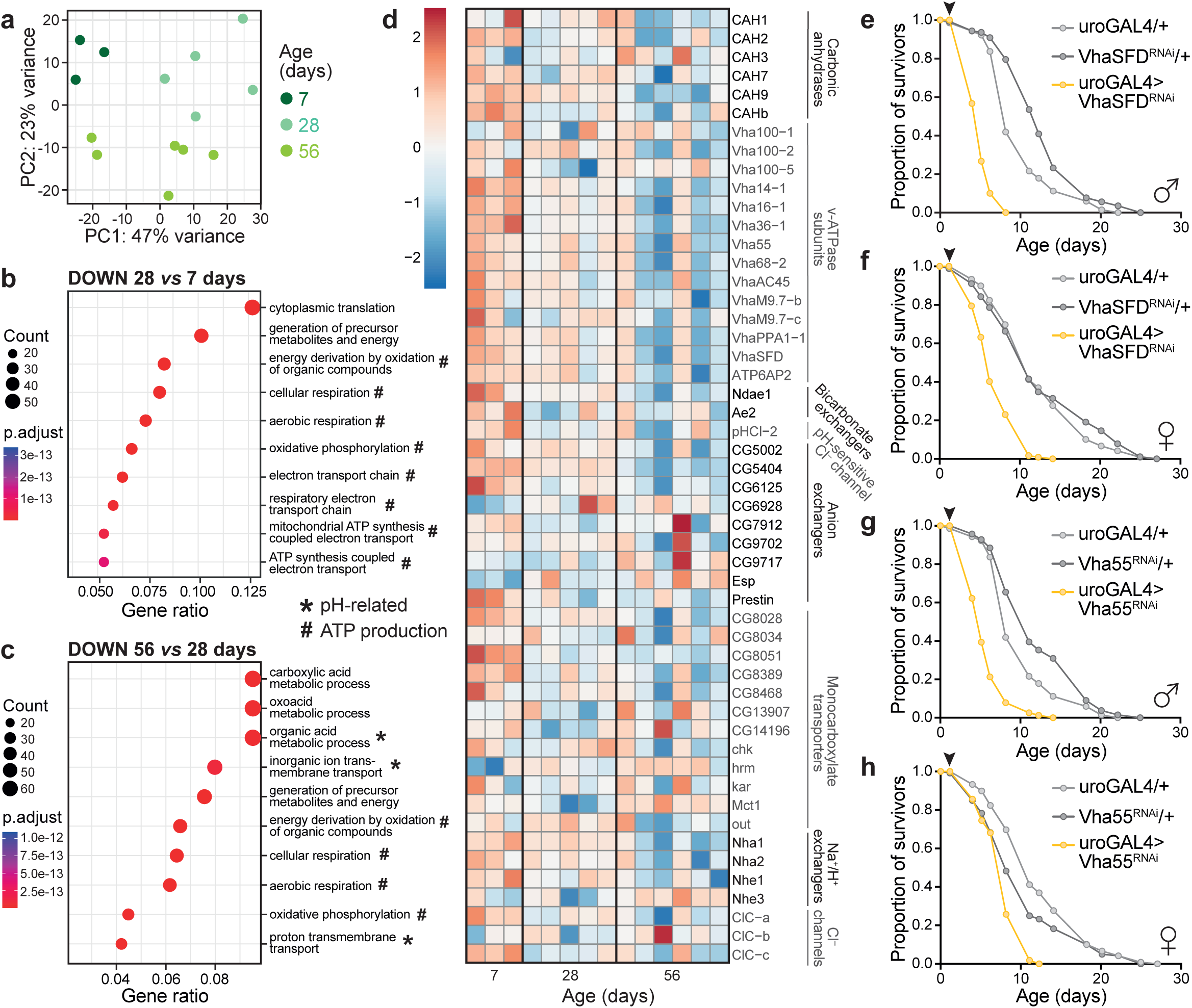
pH-related genes are downregulated in the fly excretory tract with age. **a,** Principal component analysis (PCA) of gene expression changes in dissected excretory tracts from young (7d), middle-aged (28d) and old (56d) WT (*w^Dah^*) flies. n=3-6 samples per condition. **b-c,** Bubble plots showing the top 10 gene ontology (GO) terms of downregulated genes in the excretory tract of male WT (*w^Dah^*) flies with age. 28d compared to 7d (**b**), and 56d compared to 28d (**c**). GO categories related to pH regulation (*) and ATP production by aerobic respiration (#) are indicated. **d,** Heatmap of gene expression changes related to pH regulation in the excretory tract of WT (*w^Dah^*) males with age. Selection was based on FlyBase gene categories that can alter acid-base balance through buffering or strong ion regulation: carbonic anhydrases, v-ATPase subunits, bicarbonate exchangers, chloride, monocarboxylate and sodium channels. From this subset, genes specifically enriched in the male excretory tract were included (enrichment >0.5 in either the Malpighian tubules, hindgut or rectal pad, according to FlyAtlas2)^48^. **e-h,** Survival curves of male and female flies exposed to 2.5% EG throughout their lifespan (from day 2 of adulthood, arrowheads), where v-ATPase subunit knockdown by RNAi was driven specifically in the Malpighian tubule, using the uroGAL4 driver. Tubule-specific RNAi of VhaSFD in males (**e**, n=120-142 per condition), and females (**f**, n=89-119 per condition). Tubule-specific RNAi of Vha55 in males (**g**, n=127-141 per condition), and females (**h**, n=111-120 per condition). Survival data in **e**-**h** were analyzed by Log-Rank test (see Supplementary Table 1 for full details of n numbers and p values).

From middle- to old-age, the expression of genes involved in aerobic respiration declined further, and was accompanied by decreased expression of proton and ion transport genes (Fig. 4c and Extended Data Fig. 4d). To get an overview of how pH regulation changed during aging, we focused on the subset of genes implicated in regulating acid-base balance that are expressed in the fly excretory tract, according to the FlyAtlas2 database^48^. We selected carbonic anhydrases, v-ATPase subunits, bicarbonate exchangers, chloride, monocarboxylate and other strong anion channels, and sodium channels. Strikingly, the majority of these genes were downregulated with age in the fly excretory tract (Fig. 4d), including the orthologs of genes implicated in human renal tubular acidosis (*CAH1*, *Vha55*, *Vha100-1/2/5*, *Ndae1/Ae2*)^49^. These expression changes explain the physiological loss of excretory capacity with age. To functionally validate that downregulation of these genes leads to a lower capacity to excrete acid, we knocked down the expression of several v-ATPase subunits specifically in the fly tubules and exposed the flies to acidotic stress. Both male and females flies with tubule-specific RNAi of v-ATPase subunits were indeed short-lived under acidotic stress (Figs. 4e-h and Extended Data Figs. 4e-h). Therefore, downregulation of these genes with age is sufficient to limit acid excretion, and promote systemic acidification.

### Lymph pH acidification and renal expression of acidosis-related genes in mice

To test whether our observations in flies were conserved in mammals, we examined if interstitial fluid acidification also occurred in mice. Interstitial fluid pH is understudied and may be organ-specific^50,51^. Due to the technical inaccessibility of interstitial fluids, we collected lymph from the efferent mesenteric lymph duct of mice, as previously reported^52,53^. The mesenteric lymph duct collects lymph draining the intestine and mesentery, and comprises 50% of total lymph flow in the body^52,53^. We analyzed the pH of mesenteric lymph from mice at different ages (∼10, 30 and 39 weeks) and found an age-dependent acidification (Fig. 5a) to a similar degree as in flies.

**Fig 5.**
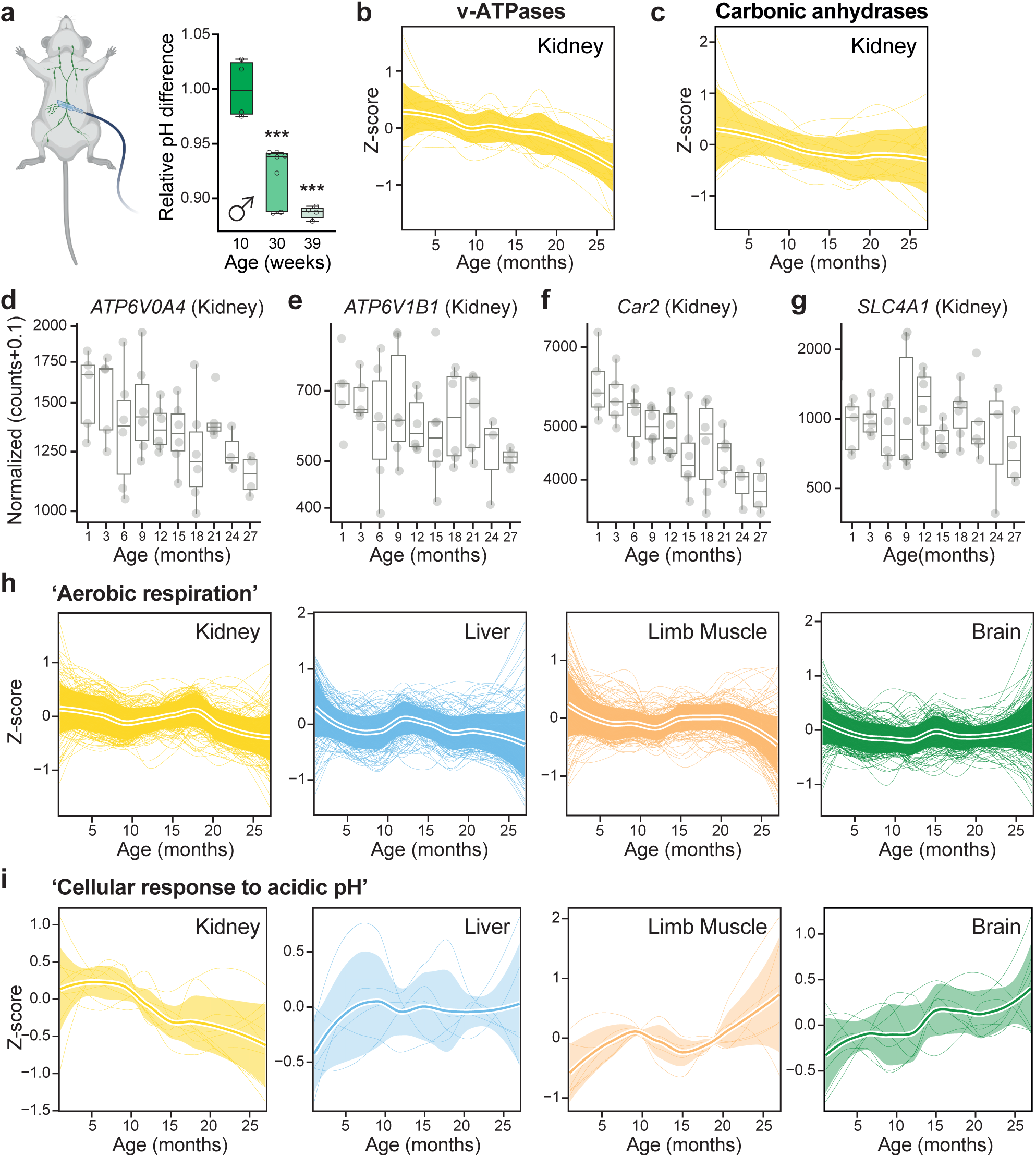
Decline in lymph pH and renal expression of acid-base genes in mice with age. **a,** Mesenteric lymph pH differences with age relative to 10-week-old male mice. Data are box-plots, with median and min-max error bars, analyzed by one-way ANOVA (Tukey) (n=4-7 replicates per condition; ***, p<0.001). **b-c,** z-Transformed gene expression trajectories of v-ATPase subunits (**b**) and carbonic anhydrases (**c**) in the mouse kidney with age (C57BL/6JN males and females)^54,55^. **d-g,** Box-plots of gene expression changes (log scale) in the kidney of C57BL/6JN mice with age^54,55^. Orthologs for pathogenic variants known to cause renal tubular acidosis in humans are displayed: *ATP6V0A4* (**d**), *ATP6V1B1* (**e**), *Car2* (**f**), and *SLC4A1* (**g**). **h-i,** z-Transformed gene expression trajectories with age in C57BL/6JN male and female mice^54,55^. Genes with the GO category terms ‘Aerobic respiration’ (**h**) and ‘Cellular response to acidic pH’^56^ (**i**) are shown for the kidney and other representative organs – liver, skeletal (limb) muscle, and brain^56^.

To explore whether insufficient acid elimination contributes to systemic acidification in mice, we analyzed the Mouse Aging Cell Atlas, which provides data on gene expression changes with age across mouse organs^54,55^. We found that all v-ATPase subunits and most carbonic anhydrases were downregulated with age in the mouse kidney (Figs. 5b,c). Interestingly, this downward trajectory does not occur universally across all tissues (Extended Data Figs. 5a,b), highlighting a particular vulnerability of the mammalian renal system to regulate pH changes with age. Next, we focused specifically on genes whose pathogenic loss-of-function variants are known to cause renal tubular acidosis in humans: the v-ATPase subunits *ATP6V0a4* and *ATP6V1B1*, the carbonic anhydrase *CA2*, and the bicarbonate exchanger *SLC4A1*^49^. Strikingly, similarly to our RNA-seq analysis of the fly excretory tract, these genes were also downregulated with age in the mouse kidney (Fig. 5d-g). Expression of *ATP6V0a4*, *ATP6V1B1* and *Car2* showed downward trends in both males and females (Extended Data Figs. 5c-e), while *SLC4A1* expression was more variable across ages (Extended Data Fig. 5f).

To complement these findings, we explored whether downregulation of genes related to ATP production would also occur in mice with age. Indeed, genes within the ontology term ‘aerobic respiration’^56^ tended to collectively decrease in the kidney with age (Fig. 5h), with similar trends in other tissues (Extended Data Fig. 5g). Interestingly, genes associated with ‘cellular response to acidic pH’^56^ were downregulated specifically in the kidney (Fig. 5i), while in most tissues their expression tended to increase with age (Extended Data Fig. 5h). These findings suggest that mouse kidneys have lower renal expression of pH-regulating and ATP production-related genes with age, which promotes systemic acidification, upregulating compensatory responses in other tissues.

## Discussion

Maintenance of extracellular pH is essential for normal tissue function. Human blood pH is regulated in a narrow range, but subclinical acidification occurs with age^12^. In interstitial fluids, where pH can have more dynamic fluctuations that escape systemic compensation, acid-base balance *in vivo* is underexplored. Using fly and mouse models, we characterized interstitial pH changes with age, and linked pH to survival. A previous study hinted at systemic acidification using diluted whole fly homogenates^29^, with limited accuracy and lacking compartment-specific resolution. To refine this further, and focus specifically on the extracellular fluid, we measured fly hemolymph pH using a cell-impermeable pH-sensitive dye. We found that fly hemolymph acidifies with age in both males and females. Although aging has sex-specific manifestations^57^, the fundamental processes that drive these changes are likely sex-independent given the ubiquitous nature of aging. Importantly, the rate of extracellular acidification strongly correlated with mortality rate across conditions, suggesting it is a common pathophysiological process preceding death. In agreement with these findings, mouse lymph acidifies with age to a similar degree, showing that age-related extracellular acidification occurs in mammals.

To explore the link between acidosis and lifespan, we established a fly model of metabolic acidosis based on EG exposure. EG-induced acidosis shortened survival in a dose-dependent manner, and flies were more susceptible to die from acidotic stress at older ages. Conversely, alkalotic treatment extended fly lifespan. In other research models, experimental manipulation of extracellular pH also alters mortality. For example, *klotho* knockout mice develop metabolic acidosis, associated with growth defects, accelerated aging-like phenotypes and shortened lifespan^58^. Treating these mice with NaHCO_3_ restores acid-base balance and, despite not preventing growth defects, extends their survival^59^. Conversely, overexpression of *klotho* in mice prolongs lifespan, independently of sex^60^. In yeast, chronological lifespan is restricted by medium acidification caused by production of acetic acid over time. Increasing the medium pH more than doubles yeast lifespan^61^. Overall, these findings indicate that extracellular acidification is a hallmark of aging across species.

Our results show that age-related systemic acidification occurs through insufficient acid elimination. The tubular portion of human kidneys changes structurally and loses function over time^62^, and can likewise contribute to chronic acidification in old age^63,64^. Interestingly, chronic acidosis in advanced chronic kidney disease and acute acidosis in critical care settings are both directly correlated with higher human mortality^65–73^. Altogether, this suggests that loss of acid elimination capacity is a conserved mechanism for insufficient response to acidosis with age, and is directly associated with mortality.

Similarly, higher dietary acid load is linked to lower renal function and increased human all-cause mortality^63,74–80^. Net endogenous acid production is determined by diet composition, including phosphate load and the metabolism of sulphur-containing amino acids, such as methionine^81^. Methionine restriction extends the lifespan of multiple species, including flies^82–85^. Supplementing ∼2 times the daily recommended intake of methionine significantly decreases urine pH in humans, and has been used as a clinical strategy to prevent kidney stones that form under alkaline conditions^86,87^. Given our findings, the increased acid load associated with excess methionine consumption may contribute to its detrimental effect on health. Indeed, even short-term consumption of a diet with lower acid load leads to systemic metabolic improvements in humans^88^. Similarly, diets with lower phosphate content may preserve renal function for longer to maintain pH balance and alter mortality^89,90^. Indeed, sevelamer, a phosphate-lowering drug, extends *Drosophila* lifespan^91^ and decreases all-cause mortality rate in patients with advanced chronic kidney disease^92,93^.

Limited capacity to regulate acid-base balance with age in multicellular organisms may allow pH fluctuations in interstitial fluids, caused by loss of perfusion, infection or bursts in acid production, to remain uncompensated for longer. Depending on the severity, repeated pH fluctuations over the lifespan may induce local irreversible damaging modifications and loss of organ function. For example, extracellular acidity can promote genomic instability, often regarded as a driver of aging^5^. Mammalian cells cultured in lower pH media (∼6.5-7.2 range compared to ∼7.4-7.8) have decreased capacity to repair DNA damage induced by independent methods^94–97^. In specific contexts, acidosis has also been associated with other hallmarks of aging, such as loss of proteostasis^98–100^, inhibition of macroautophagy^101^, increased inflammation with impaired innate and adaptive immune function^102^, and altered nutrient signalling and mitochondrial dysfunction^103,104^. Senescent cells have lower intracellular pH compared to proliferating cells^105,106^, which can be induced by lysosomal membrane damage^106^, but may also be promoted by an acidic extracellular environment^107^. Therefore, acid-base imbalance may precede and underlie other established aging hallmarks. These mechanistic interactions warrant further exploration in the pathophysiology of age-related diseases and geriatric syndromes.

Overall, our study sheds light on extracellular pH regulation in aging, and suggests acidification as a fundamental evolutionarily conserved pathophysiological process that increases disease risk with age to ultimately limit lifespan. Future research should aim to dissect the role of interstitial fluid pH fluctuations in tissue function, and develop novel strategies to maintain pH homeostasis systemically and across organs with age.

## Methods

### Fly maintenance

Flies were kept in an incubator or room with controlled temperature and humidity (65%) on a 12 h:12 h light-dark cycle. Experiments were performed at 25°C unless otherwise stated. Parental flies were put into cages with apple juice-agar plates supplemented with live yeast paste. After 24 h, eggs were collected by rinsing the plates with phosphate-buffered saline (PBS, Sigma 524650). Flies were reared at constant density by pipetting 16 µL of egg suspension using a wide-bore 200 µL tip into 200 mL bottles with SYA medium. Newly eclosed adults were transferred onto bottles of fresh medium without anesthesia and kept as a mixed population for 2 days to allow maturation and mating. Flies were then sorted using a light microscope by sex and genotype under mild CO_2_ anesthesia into experimental vials (∼15-20 flies per vial). Unmated females were selected on ice following eclosion and transferred to bottles before being sorted into vials after 2 days. Experiments were performed on mated males and females, unless otherwise stated.

### Fly media and drug supplementation

Flies were kept on SYA medium, unless otherwise stated. SYA consists of 5% (w/v) sucrose (granulated sugar), 10% (w/v) Brewer’s yeast (903312, MP Biomedicals) and 1.5% (w/v) agar (A7002, Sigma), supplemented with 30 mL/L of 10% (w/v) nipagin (H5501, Sigma; prepared as a solution in 95% ethanol) and 0.3% (v/v) propionic acid (P1386, Sigma) as preservatives, added once the food had cooled down to <60°C. Apple juice-agar plates for cages were prepared with 25% (v/v) apple juice, 2.3% (w/v) glucose, 2.3% (w/v) agar and 1.5% (v/v) nipagin. The chemically-defined (holidic) medium with an amino acid profile similar to yeast was prepared as previously described^108^. Added drugs or chemicals were supplemented to standard media once cooled down to <60°C, including pH adjustments with 10 M NaOH. Medium pH was measured using a food pH meter (PH60S, Apera Instruments). Ethylene glycol (324558, Sigma) was added directly to the food. 100 mM NaHCO_3_ (S5761, Sigma), 10-100 mM Tris base (T6066, Sigma) and 1 mM LiCl (L9650, Sigma) were added to the food dissolved in deionized water. Streptomycin (S6501, Sigma) and 50 μM rapamycin (R-5000, LC Laboratories) were dissolved in 100% ethanol. 15.6 μM trametinib (T-8123, LC Laboratories) was dissolved in dimethyl sulfoxide (472301, Sigma). Equivalent volumes and concentrations of vehicles were added to control SYA medium. Drug treatments were started at day 2 of adulthood unless otherwise stated.

### Fly strains and genotypes

The *white Dahomey* (*w^Dah^*) strain of *Drosophila melanogaster* was used as the WT background for most experiments. The isogenic control line *w^1118^* and the WT strain *Ore^R^*were occasionally used as specified. The *w^Dah^*, *w^1118^* and Δ*dilp2-3,5* flies were originally gifted by Linda Partridge, University College London. The *Ore^R^* flies were kindly provided by Irene Miguel-Aliaga, MRC LMS. The following UAS-RNAi lines were obtained from the Vienna *Drosophila* Resource Center (VDRC, www.vdrc.at):

UAS-VhaSFD^RNAi^ VDRC #47471: RRID:Flybase_FBst0467325 (*w^1118^*;P{GD8795}v47471)

UAS-Vha55^RNAi^ VDRC #46553: RRID:Flybase_FBst0466765 (*w^1118^*;;P{GD9363}v46553/TM3)

UAS-Vha13^RNAi^ VDRC #25985: RRID:Flybase_FBst0456166 (*w^1118^*;P{GD10564}v25985)

UAS-Vha44^RNAi^ VDRC #46563: RRID:Flybase_FBst0466770 (*w^1118^*;;P{GD10617}v46563).

The tubule-specific driver line uroGAL4 (*w^1118^*, uroGAL4;;)^109^ was gratefully received from Julian Dow, University of Glasgow. The UAS-RNAi lines and uroGAL4 were backcrossed into the *w^1118^* background.

### Fly survival

Survival experiments were set up as described above, typically with ∼140-150 flies per condition (∼7-8 vials of 20 males or ∼10 vials of 15 females). Deaths and censors were scored every 2-3 days, as the flies were transferred into vials with fresh media. Under stress conditions, deaths were scored more frequently, up to twice a day. During the transfer, dead flies were removed from vials^110^.

### Fly feeding behavior

Fly feeding was assessed by observation of proboscis extension. 5 flies were set up per vial and acclimatized to the temperature and humidity-controlled room before scoring. Feeding behavior was directly observed and recorded over 1 h in cycles of approximately 5 min. Experimental conditions were blinded and only revealed at the end for data analysis.

### *In vivo* fly respirometry

A high-resolution insect respirometry system developed by Sable Systems was set up as previously described^111,112^. Groups of 3 flies were transferred into respirometry chambers without anesthesia. Each chamber had a food source using a loose plastic cuvette cap as a container. The reference chamber (with no flies) had the same food source in it. Changes in gas (oxygen, carbon dioxide and water vapour) partial pressures were measured in cycles of 48 min. Peaks from the baseline gas partial pressure were analyzed as the area under the curve. Data were typically recorded over 24 h using the ExpeData software provided by Sable Systems. The same software was used to analyze the data, with a custom macro kindly provided by Andreas Mölich, Sable Systems Europe GmbH.

In summary, the analysis consists of the following steps: calculation of mean flow-rate (FR); correction of CO_2_ and O_2_ channels for lag; calculation of fractional concentration of CO_2_ from ppm and multiplication with FR; calculation of fractional concentration of O_2_ from % and multiplication with FR; conversion of both to µl/min, and division by cycle length. For CO_2_, drift is corrected (set to zero) and possible artefacts are removed (despike). RQ is calculated by dividing CO_2_ values by the O_2_.

### Fly egg bleaching

Eggs were bleached after collection from apple juice-agar plates with PBS into a cell strainer. Eggs were immersed and gently swirled in thick bleach for 2 min, and then rinsed thoroughly with water before being transferred to SYA medium as above.

### Fly bacterial load quantification

Fly bacterial load was quantified by culturing whole fly homogenates, as previously described^38^. Briefly, three 7-day-old flies were placed into 1.5 mL Eppendorf tubes and homogenized using a sterile plastic pestle. 100 µM of homogenate was pipetted onto petri dishes containing MRS agar (CM1153B, Thermo Scientific), selective for *Lactobacilli*, and distributed using a sterile spreader. Plates were left to dry before being closed and sealed with parafilm, and incubated at 25°C. Bacterial colonies were counted manually after 72 h.

### Adult fly hemolymph collection

To extract hemolymph from adult flies, live cohorts of 12 females or 15 males were decapitated under ice anesthesia using a scalpel blade. The bodies were placed upside down into cut 200 µL pipette tips within a 1.5 mL tube and gently centrifuged for 15 min at 1,500 x g, 4°C to drain the hemolymph (obtaining ∼1 µL per sample). The tube was then snap-frozen in liquid nitrogen and kept on dry ice during the collection process before being stored at −80°C until required. Samples which were not clear (i.e. cloudy or containing eggs) were discarded.

### Mouse maintenance and lymph collection

C57BL6/J mice (6-8 weeks old) were randomized and housed in groups of 2-5 on a 12 h:12 h light/dark cycle at 22-25°C and with 35-37% humidity. Animals were fed a semi-purified normal chow diet (102119, Barastoc) for 3, 23 or 32 weeks. Following this feeding period, mice were anesthetized and the efferent mesenteric lymphatic duct was cannulated, as described previously ^52,53^. Lymph was collected for up to 4 h in the morning in non-fasted animals. Animal work was approved by the Monash Institute of Pharmaceutical Sciences Animal Ethics Committee and conducted according to Australian National Health and Medical Research Council (NHMRC) guidelines for the care and use of animals in research.

### Hemolymph and lymph pH analysis

Hemolymph pH was measured based on the spectrophotometric profile of the pH-sensitive cell-impermeable pyranine dye, adapted from a previous report for larval hemolymph^17^. In short, to obtain the pH of a hemolymph sample, 1 µL of the pH-sensitive dye pyranine (2.4 mM in water; H1529, Sigma) was mixed with 1 µL of hemolymph. The absorbance ratio of the pyranine protonated and deprotonated peaks was measured by spectrophotometry using a Nanodrop One instrument (Thermo Scientific). These peaks were respectively at 405 and 450 nm with automated baseline correction (or 425 and 484 nm without) (Extended Data Figs. 1a,c). To obtain a standard curve, 1 µL of pyranine dye was mixed with 1 µL of 50 mM Tris-HCl solutions at different pHs, ranging between 6.6 and 7.8. Values were plotted as a second-degree polynomial equation with pH on the y-axis and the absorption ratio on the x-axis (Extended Data Fig. 1c). We confirmed that the pH of 3^rd^ instar larvae hemolymph was ∼7.4-7.5, as reported previously^17^, while hemolymph pH of young (7d) flies was ∼7.1-7.2. Mouse lymph pH was measured using the same approach, mixing 20 µL of lymph with 80 µL of pyranine dye (100 nM). Lymph pH was ∼8.3 in young (∼10 week-old) mice. pH values are presented as relative differences to the control condition from calculated pH based on the Tris-HCl standard curve.

### Fly excreta pH analysis

Groups of 5 flies were transferred without anesthesia to a Petri dish containing a 1/4 wedge of SYA medium supplemented with the pH-sensitive dyes bromophenol blue (0.5% w/v; B0126, Sigma) or bromocresol purple (0.5% w/v; 114375, Sigma)^42^. SYA medium was prepared without propionic acid and pH was adjusted to 4.0 or 5.6, respectively. The flies were kept in the dishes for 24 h at 25°C. The plate lids were scanned and the excreta were analyzed using T.U.R.D. software (v0.8)^43^, with the following settings: block size 25, offset 5, minimum size 35, maximum size 3500, shape circle, size 3, and circularity threshold 0.70. Excreta with hue above 300 (end of blue) and below 30 (end of yellow) were excluded.

### Malpighian tubules deposit scoring

Flies were anesthetized on ice and dissected in chilled PBS under a light microscope. Deposits were scored according to the proportion of stones within each tubule, as previously described^26^, with grading from 0 to 4: ‘0’, ∼clear tubule (<5% stones); ‘1’, <25% stones; ‘2’, 25-50% stones; ‘3’, 50-75% stones; ‘4’, >75% stones. The score per fly was averaged from its four tubules.

### Excretory tract RNA extraction

Flies were anesthetized on ice and the excretory tract (Malpighian tubules, hindgut and rectal ampulla) was dissected by pulling their intestine from the rectal pad in an RNA stabilization solution (RNAlater; AM7020, Invitrogen). The intestine was cut above the insertion of the Malpighian tubules, and the dissected tissue was transferred into 2 mL RNase-free tubes with beads on dry ice (15 samples per tube). Frozen samples were stored at −80°C. Before RNA extraction, RNAlater solution was removed after centrifugation (16,000 rpm, 5 min, 4°C) and tissues were homogenized in RLT buffer (30 s, 6000 rpm, Precellys homogenizer). Total RNA was extracted following the protocol of the RNeasy Mini kit (74104, Qiagen).

### RNA processing and sequencing

RNA quality and concentration was assessed using the Agilent 2100 Bioanalyzer RNA 6000 Nano assay. polyA enrichment was performed on 200 ng total RNA, using the NEBNext® Poly(A) mRNA Magnetic Isolation module. NGS RNA-seq libraries were generated using the NEBNext® Ultra™ II Directional RNA Library Prep Kit for Illumina® according to manufacturer’s instructions. Library quality was assessed using the Agilent 2100 Bioanalyzer High-Sensitivity DNA assay. For each sample, a minimum of 20 million Paired End 60bp reads (dual 8bp indexing) were generated on an Illumina NextSeq 2000.

### RNA-seq analysis

RNA-seq reads were processed using cutadapt (v4.1)^113^ to remove Illumina adapters, to quality trim at Q20, and to filter read pairs containing N bases or where either read <31 bps. Processed reads were quantified with Salmon (v1.9.0)^114^ using transcripts from the Dmel 6.48 Ensembl annotations. Salmon’s expectation maximization procedures were set to enable modelling of sequencing and GC biases. Within R, transcript level counts from Salmon were imported and aggregated to the gene level using tximport (v1.30.0)^115^. Normalization and differential expression analyses were performed with DESeq2 (v1.40.2)^116^. Independent hypothesis weighting was conducted to optimize the power of p-value filtering^117^ and log2 fold change shrinkage was performed using ashr^118^ to reduce the impact of low expression on estimation of fold change values. Multiple correction testing adjustments were optimized by passing thresholds for both false discovery rate and log2 fold change at the point of results generation within DESeq2. Genes passing significance thresholds were used to assess for enrichment of GO (BP: Biological Process) gene sets using clusterProfiler (v4.8.1)^119^.

### Statistics and reproducibility

Survival curves were plotted in Microsoft Excel and analyzed by Log-Rank test using a custom macro (see Supplementary Tables 1 and 2 for full details of n numbers and p values). All other data were plotted using GraphPad Prism (v10) software and analyzed either by unpaired two-tailed *t*-test or one-way ANOVA with Tukey correction for multiple comparison, as appropriate. Significance of probability values: non-significant (ns), p>0.05; *, p<0.05; **, p<0.01; ***, p<0.001. No statistical methods were used to predetermine sample sizes, but our n numbers are similar to those reported in previous studies)^26^. Data distribution was assumed to be normal, but this was not formally tested. Flies of the same genotype were randomly assigned to treatments/food conditions during experimental set-up. Feeding assays were scored blind. Other assays (e.g. lifespans, stress assays, hemolymph analysis) were performed unblinded, due to the requirement of the investigator to document genotypes and food conditions.

## Data availability

All data that support the findings of this study are available from the corresponding authors upon reasonable request. Full details for all survival data (n numbers, p values) are provided in Supplementary Tables 1 and 2. The RNA-seq dataset has been deposited in the GEO repository (accession: GSE263182).

## Acknowledgements

We acknowledge the support of Laurence Game and the LMS Genomics Facility in processing RNA-seq samples. We acknowledge use of the Jex HPC cluster^120^, and the resources provided by the LMS IT and Bioinformatics Facilities in analyzing the RNA-seq data. We thank all current and former members of the Cochemé group for their input, in particular Claudia Lennicke and Marcela Buricova. We are grateful to Filipe Cabreiro and Susumu Hirabayashi for valuable feedback. Figures 1a, 3a, 5a, S1a and S2b were created with BioRender.com. For the purpose of open access, the author has applied a Creative Commons Attribution (CC BY) licence to any Author Accepted Manuscript version arising.

## Funding

This work was funded by the Medical Research Council (MC-A654-5QB90) to H.M.C., New Zealand Health Research Council Program Grant (21-714) to N.L.T., Australian National Health and Medical Research Council (NHMRC) Investigator Grant (#2017903) to E.C., and Monash University International Scholarship to Y.X. E.d.S. was supported by an ERDA award from the Institute of Clinical Sciences, Imperial College London.

## Author contributions

E.d.S. conceived the project, supervised by H.M.C. E.d.S. designed, performed and analyzed fly experiments, with technical support from A.F. Mouse lymph samples were collected by E.C. and analyzed by Y.X., supervised by N.L.T. M.E.T. and I.A. processed the RNA-seq samples. G.Y. analyzed the RNA-seq data. E.d.S. and H.M.C. wrote the manuscript, with input from all authors.

## Competing interests

N.L.T. and E.C. are inventors of the lymph-directing glyceride prodrug technology (WO2016023082 and PCT/AU2020/050997), which has been patented and licensed via a commercial agreement with PureTechHealth and Seaport Therapeutics. The remaining authors declare no competing interests.

## Extended Data - Figure Legends

**Extended Data Fig. 1.**
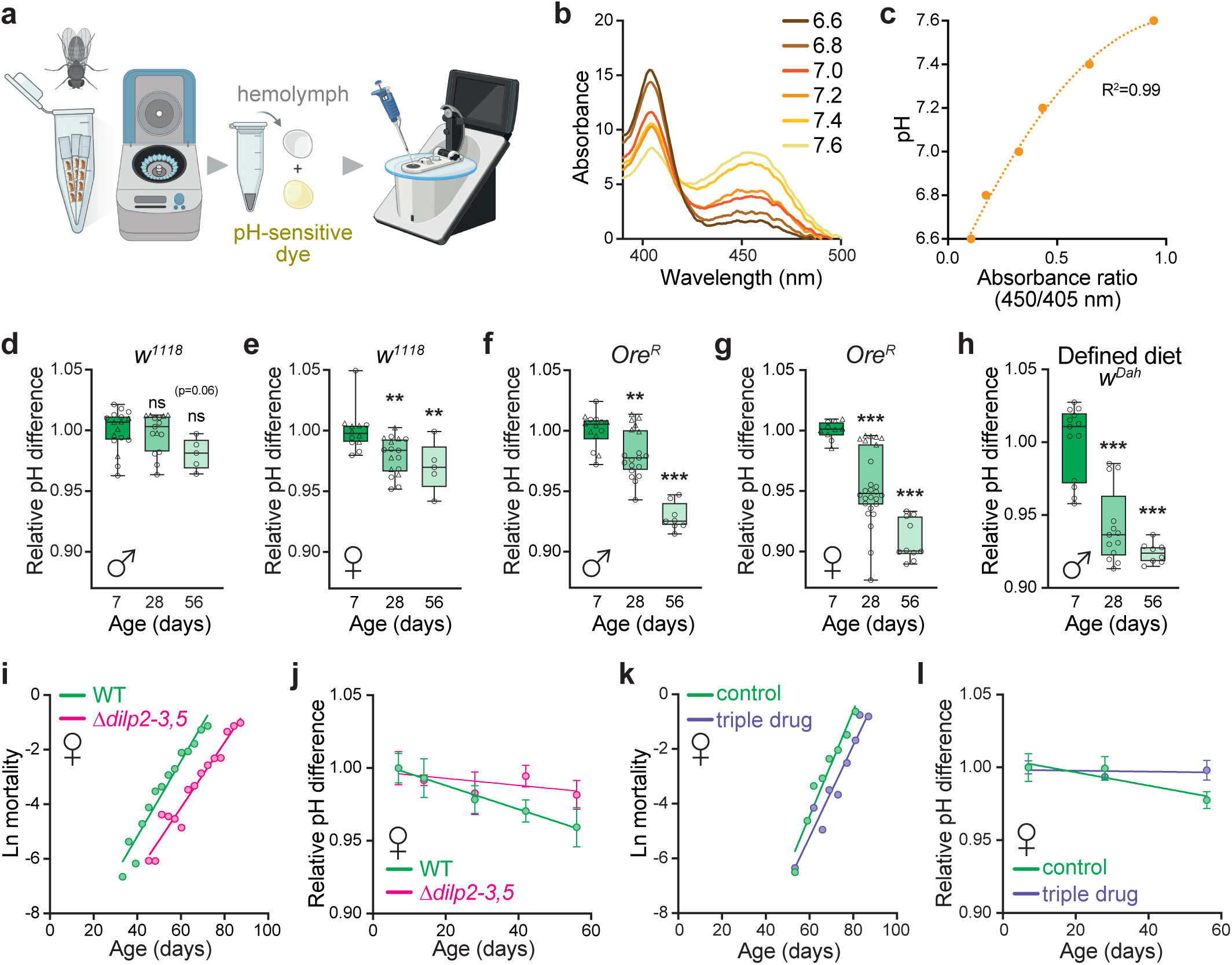
Extracellular acidification of adult *Drosophila* correlates with mortality rate. **a-c,** Assay of extracellular fluid pH. Scheme of adult fly hemolymph collection and pH measurement (**a**). Hemolymph is drained from decapitated flies by gentle centrifugation, then combined with a cell-impermeable pH-sensitive pyranine dye before spectrophotometric analysis on a NanoDrop. Absorbance spectra of pyranine dye when mixed with buffer solutions (pH 6.6 to 7.6). The protonated (405 nm) and deprotonated (450 nm) absorbance peaks of pyranine are pH-dependent (**b**). Standard curve of buffer solution pH for each 450/405 nm ratio of absorbance, fitted using a second-degree polynomial equation (**c**). **d-e,** Hemolymph pH differences with age in *w^1118^* flies relative to 7d. Males (**d**, n=5-17 replicates) and females (**e**, n=5-17 total replicates), each pooled from 2 independent experiments. **f-g,** Hemolymph pH differences with age in *Ore^R^* flies relative to 7d. Males (**f**, n=8-19 total replicates) and females (**g**, n=9-26 total replicates), each pooled from 2 independent experiments. **h,** Hemolymph pH differences with age of WT (*w^Dah^*) males raised on a chemically-defined diet, relative to 7d (n=8-13 replicates per condition). Data in **d**-**h** are box-plots, with median and min-max error bars. Statistical analysis was performed by one-way ANOVA (Tukey) against the d7 timepoint. **i,** Mortality rates of WT (*w^Dah^*) and *Δdilp2-3,5* females flies, corresponding to the survival curves in Fig. 1i. The mortality trajectories are shifted, but the slopes of their respective fitted trendlines are not statistically different (p=0.15). **j,** Rate of hemolymph acidification in WT (*w^Dah^*) and *Δdilp2-3,5* females, corresponding to the pH data in Fig. 1j. Data are means ±SD (changes are relative to the 7d condition of each genotype). **k,** Mortality rates of WT (*w^Dah^*) females treated with a triple drug combination of 50 μM rapamycin, 15.6 μM trametinib and 1 mM lithium, corresponding to the survival curves in Fig. 1k (slopes, p=0.38). **l,** Rate of hemolymph acidification in WT (*w^Dah^*) females in response to triple drug treatment, corresponding to the pH data in Fig. 1l. Data are means ±SD, relative to the 7d condition. Slopes of fitted trendlines are significantly different (p<0.01). Each pH data point is based on the hemolymph collected from a cohort of 15 male or 12 female flies, with different symbols corresponding to independent experiments. Slopes of mortality (**i**,**k**) and relative pH acidification rates with age (**j**,**l**) were calculated by simple linear regression and compared by analysis of covariance. ns, p>0.05; **, p<0.01; ***, p<0.001.

**Extended Data Fig. 2.**
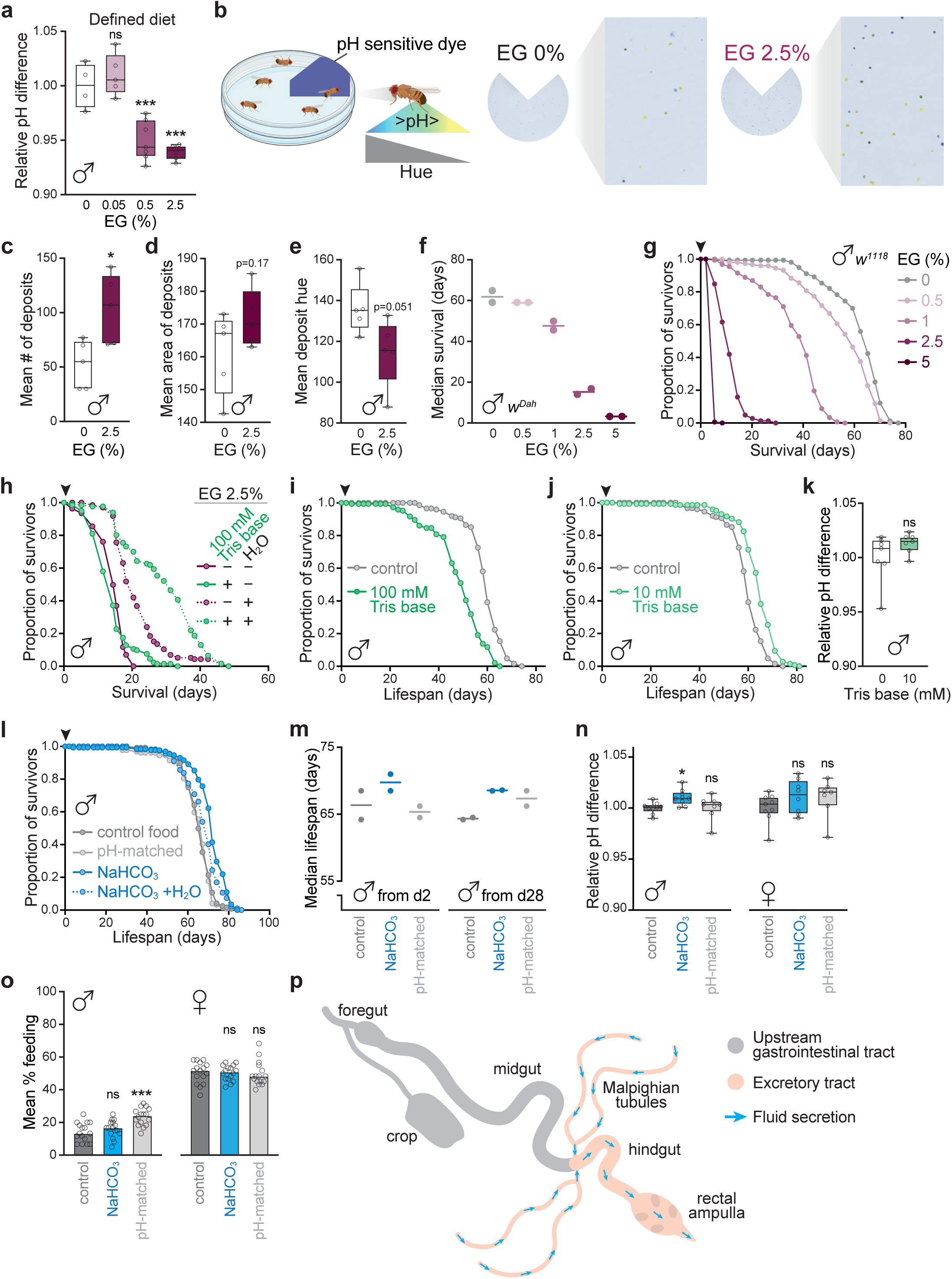
Susceptibility to acidotic stress increases with age. **a,** Hemolymph pH differences in male WT (*w^Dah^*) flies after 7 days of exposure to EG (0.05-2.5%) in chemically-defined diet, relative to untreated controls (n=4-7 replicates per condition). **b,** Scheme of adult fly excreta pH analysis. Flies are transferred to Petri dishes with a wedge of food containing the pH-sensitive dye bromocresol purple. After 24 h, the plate lids are scanned and the excreta are analyzed by number, area and hue. Lower hue (towards yellow) corresponds to more acidic deposits. Representative images of excreta from young (7d) male flies exposed to 2.5% EG compared to untreated controls. **c-e,** Analysis of excreta from young WT males exposed to 2.5% EG for 7 days compared to untreated controls. Mean number of deposits (**c**), mean area of deposits (**d**), and mean deposit hue (**e**) per plate (n=5 plates, each with 5 flies). **f,** Summary of median survival data for WT (*w^Dah^*) males exposed to EG (0.5-5%) throughout their lifespan, from day 2 of adulthood. Data are means of 2 independent repeats (see Fig. 2b). **g,** Survival curves of male *w^1118^* flies exposed to EG (0.5-5%) throughout their lifespan, from day 2 of adulthood (arrowhead). n=152-160 flies per condition. **h,** Survival curves of male WT (*w^Dah^*) flies exposed to 2.5% EG throughout their lifespan, from day 2 of adulthood (arrowhead), supplemented with 100 mM Tris base and/or a water source. n=133-139 flies per condition. **i-j,** Survival curves of male WT (*w^Dah^*) flies treated with 100 mM (**i**) or 10 mM (**j**) Tris base continuously from day 2 of adulthood (arrowhead). n=151-152 flies per condition. **k,** Hemolymph pH differences of young WT (*w^Dah^*) males treated with 10 mM Tris base for 7 days, compared to untreated controls. n=6-7 replicates per condition. **l-m,** Lifespan of WT (*w^Dah^*) males treated with 100 mM NaHCO_3_ or food with the pH adjusted by NaOH to match the NaHCO_3_ pH effects. Representative survival curves of flies treated continuously from day 2 of adulthood (arrowhead), including a condition where an extra source of water was provided (**l**, n=122-157). Summary of median survival for flies treated with 100 mM NaHCO_3_ from day 2 or day 28 of adulthood (**m**, n=122-157 flies per condition). **n,** Hemolymph pH differences of young WT (*w^Dah^*) flies treated for 7 days with 100 mM NaHCO_3_ or pH-matched food (NaOH), compared to flies raised on control diet. n=7-9 replicates per condition. **o,** Mean proportion of flies feeding per vial during 60 min, by direct observation of proboscis extension. Yound male and female WT (*w^Dah^*) flies were pre-treated for 7 days with 100 mM NaHCO_3_ or pH-matched food (NaOH), and maintained on the same conditions during the experiment. Data are means of n=16 vials, each with 5 flies. **p,** Scheme of the fly gastrointestinal and excretory tract. Secreted fluids from the two pairs of Malpighian tubules join the gastrointestinal tract at the midgut-hindgut junction, and mix with digestive waste along the hindgut and rectal ampulla before being eliminated as feces. Survival data in **g**-**j** and **l-m** were analyzed by Log-Rank test (see Supplementary Table 2 for full details of n numbers and p values). Each pH data point is based on the hemolymph collected from a cohort of 15 male or 12 female flies. Graphs in **a**, **c**-**f**, **k** and **n** are box-plots, with median and min-max error bars. Statistical significance was determined by one-way ANOVA (Tukey) for **a**, **n** and **o**, or unpaired two-tailed *t*-test for **c**-**f** and **k**, against the untreated control. ns, p>0.05; *, p<0.05; ***, p<0.001.

**Extended Data Fig. 3.**
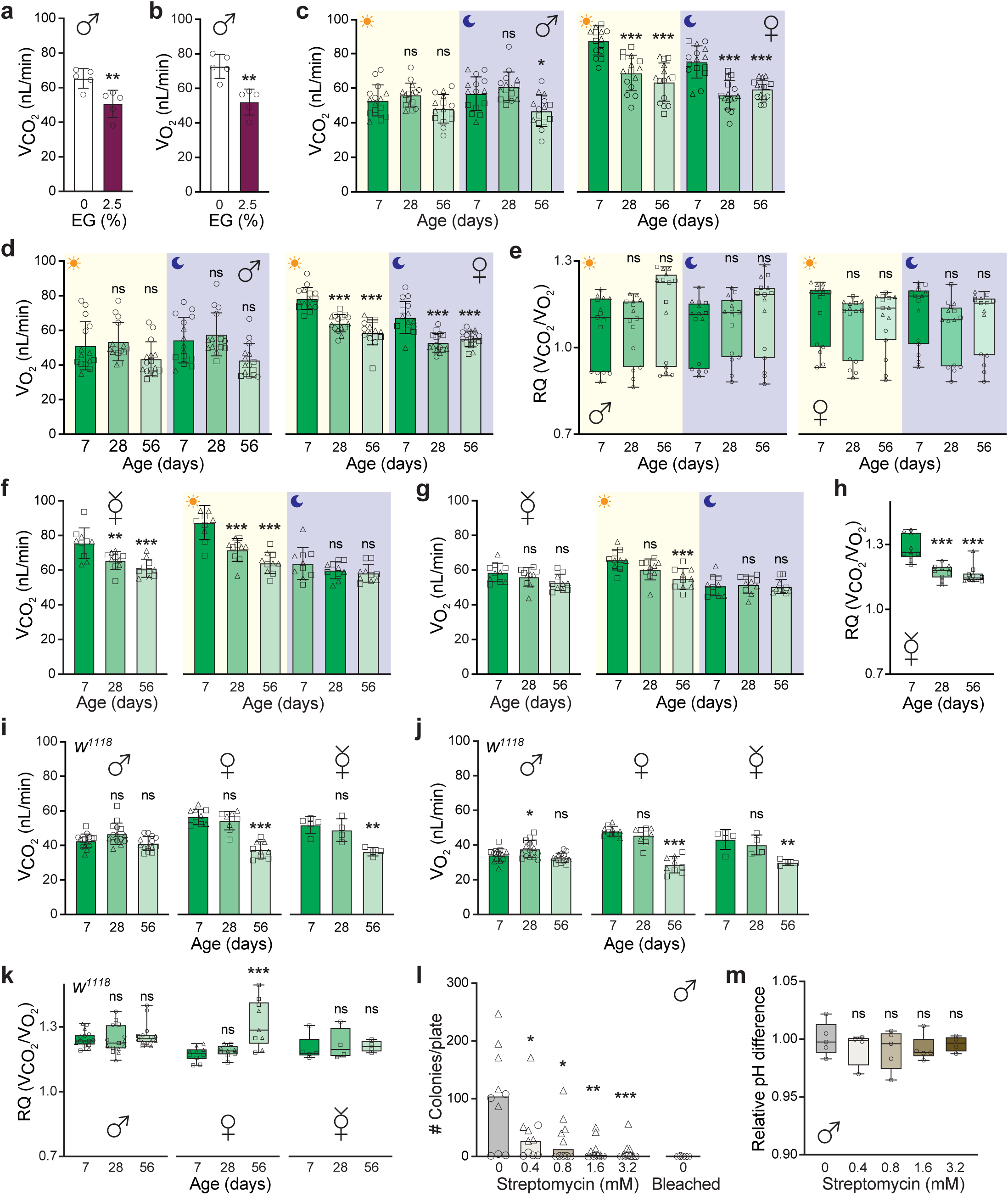
Global acid elimination declines with age in *Drosophila.* **a-b,** Respirometry analysis of young WT (*w^Dah^*) males following exposure to 0 or 2.5% EG for 7 days (n=5 chambers per condition, each with 3 flies, measured over 24 h). Means ±SD of CO_2_ production (**a**) and O_2_ consumption (**b**) for n=5 replicates, analyzed by a two-tailed unpaired *t*-test. **c-e,** Respirometry analysis of mated WT (*w^Dah^*) males and females with age, broken down by light-dark cycle, represented by the sun and moon icons respectively. Mean ±SD of CO_2_ production (**c**), mean ±SD of O_2_ consumption (**d**), and RQ (**e**). n=15 total replicates per condition (=chamber containing 3 flies), pooled from 3 independent experiments, as indicated by the different symbols. **f-h,** Respirometry analysis of unmated WT (*w^Dah^*) females with age, broken down by light-dark cycle, represented by the sun and moon icons respectively. Mean ±SD of CO_2_ production (**f**), mean ±SD of O_2_ consumption (**g**), and RQ (**h**). n=10 total replicates (chamber containing 3 flies) per condition, pooled from 2 independent experiments, as indicated by the different symbols. **i-k,** Respirometry analysis of *w^1118^* flies (mated males and females, and unmated females) at different ages. Mean ±SD of CO_2_ production (**i**), mean ±SD of O_2_ consumption (**j**), and RQ (**k**). n=15 total replicates (chamber containing 3 flies) per condition for mated males, pooled from 3 independent experiments, as indicated by the different symbols. n=10 replicates per condition for mated females, pooled from 2 independent experiments, and n=5 replicates for unmated females. **l,** Number of bacterial colonies per MRS plate with homogenates of 3 WT (*w^Dah^*) males either treated with a range of streptomycin concentrations for 7 days during adulthood (n=10 plates, pooled from 2 independent experiments), or eclosed from bleached eggs (n=5 plates). **m,** Hemolymph pH differences in WT (*w^Dah^*) males treated with streptomycin (0.4-3.2 mM) for 7 days, relative to flies raised on control food. n=4-5 replicates per condition. Each pH data point is based on the hemolymph collected from a cohort of 15 flies. Graphs in **e**, **h**, **k** and **m** are box-plots, with median and mix-max error bars. Data in **c**-**m** were analyzed by one-way ANOVA (Tukey), against the young/untreated control.

**Extended Data Fig. 4.**
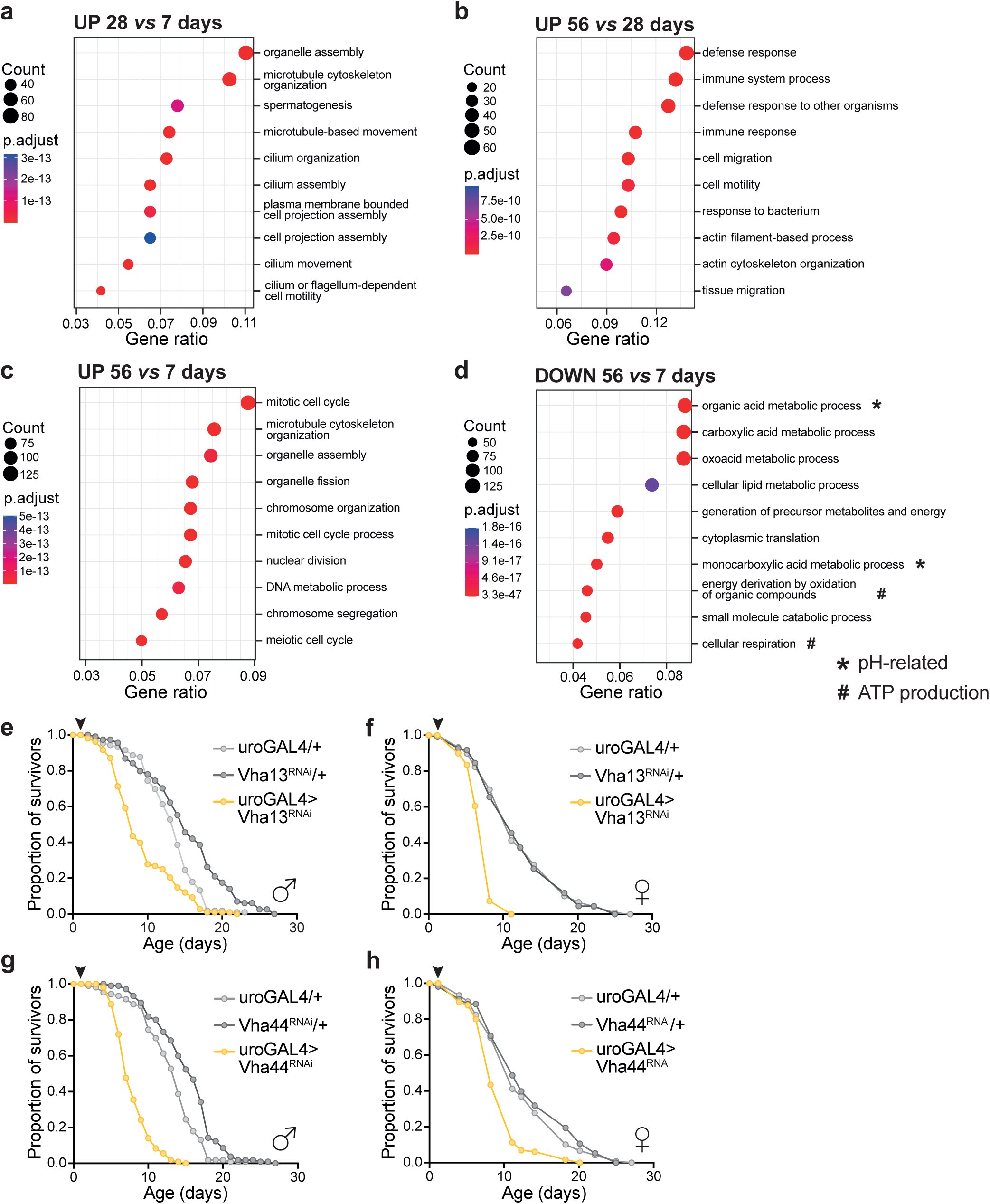
pH-related genes are downregulated in the fly excretory tract with age. **a-c,** Bubble plots listing the top 10 gene ontology (GO) terms of upregulated genes in the excretory tract of WT (*w^Dah^*) males with age: 28d compared to 7d (**a**), 56d compared to 28d (**b**), 56d compared to 7d (**c**). **d,** Bubble plot with the top 10 GO terms of downregulated genes in the excretory tract of 56d male WT (*w^Dah^*) flies with age compared to 7d. Genes related to pH regulation (*) and ATP production by aerobic respiration (#) are signed. **e-h,** Survival curves of male and female flies exposed to 2.5% EG throughout their lifespan, (from day 2 of adulthood, arrowheads), where v-ATPase subunit knockdown was driven specifically in the Malpighian tubule, using the uroGAL4 driver. Tubule-specific RNAi of Vha13 in males (**e**, n=106-114 per condition), and females (**f**, n=109-110 per condition). Tubule-specific RNAi of Vha44 in males (**g**, n=127-141 per condition), and females (**h**, n=111-120 per condition). Survival data in **e**-**h** were analyzed by Log-Rank test (see Supplementary Table 2 for full details of n numbers and p values).

**Extended Data Fig. 5.**
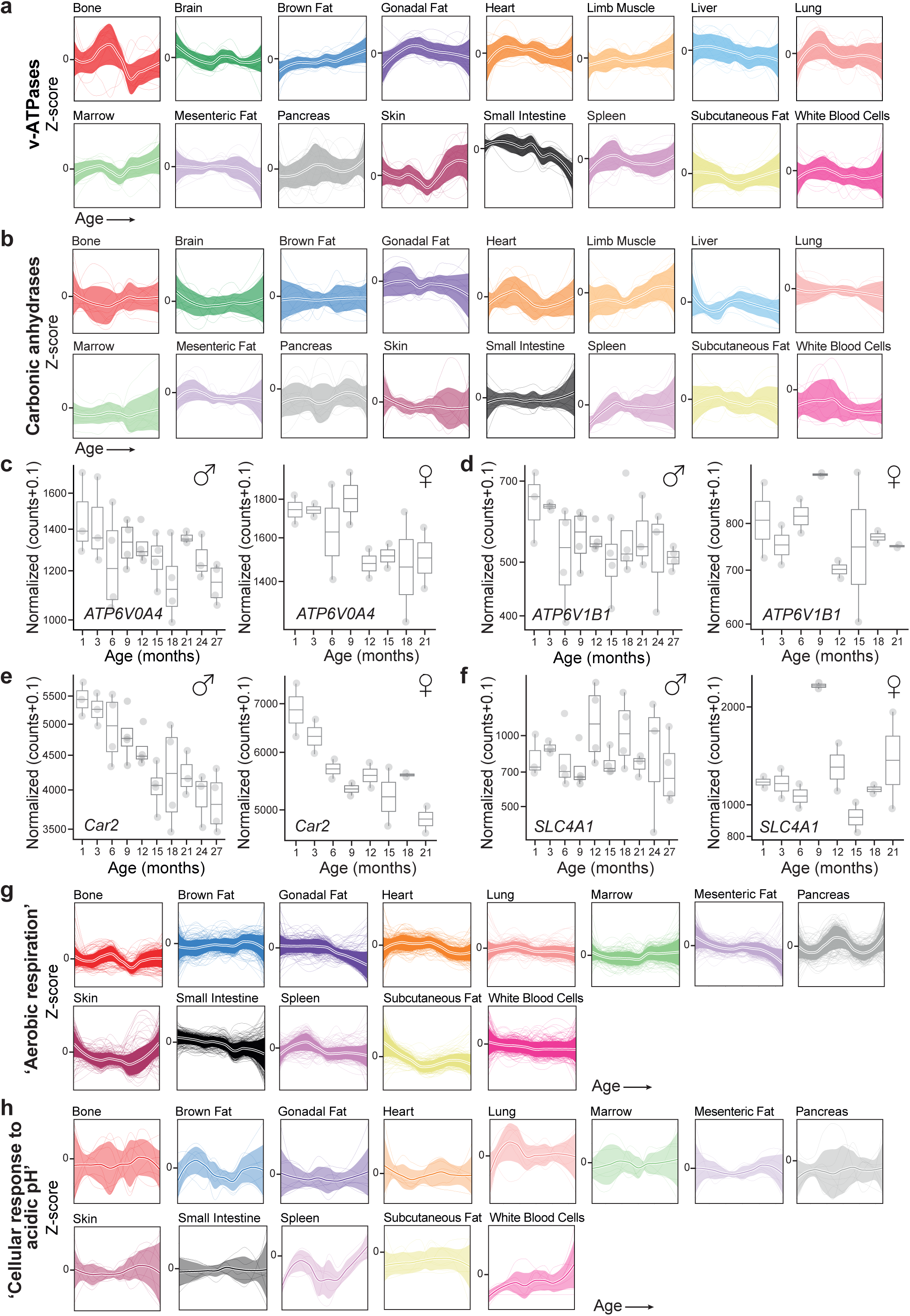
Decline in lymph pH and renal expression of acid-base genes in mice with age. **a-b,** z-Transformed gene expression trajectories of v-ATPase subunits (**a**) and carbonic anhydrases (**b**) across mouse tissues besides the kidney with age (C57BL/6JN males and females)^54,55^. **c-f,** Box-plots of gene expression changes (log scale) in the kidney of C57BL/6JN mice with age, with males and females plotted separately (see Figs. 5d-5g for pooled data)^54,55^. Orthologs for pathogenic variants known to cause renal tubular acidosis in humans are displayed: *ATP6V0A4* (**c**), *ATP6V1B1* (**d**), *Car2* (**e**), and *SLC4A1* (**f**). **g-h,** z-Transformed gene expression trajectories with age in C57BL/6JN male and female mice^54,55^. Genes with the GO category terms ‘Aerobic respiration’ (**g**) and ‘Cellular response to acidic pH’ (**h**) are shown for tissues besides the kidney, liver, skeletal muscle and brain (see Figs. 5h,i)^56^.

**Supplementary Table 1.**
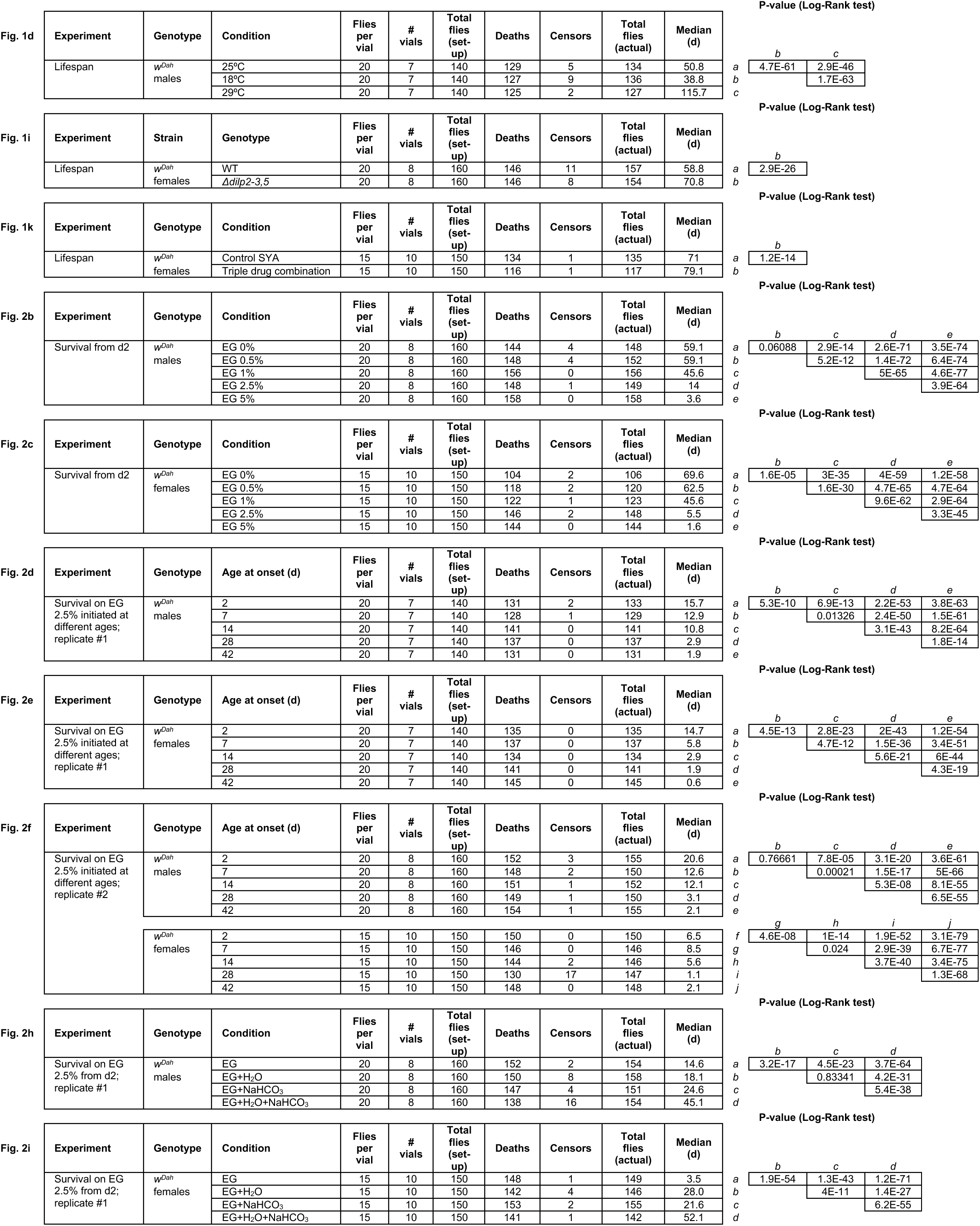
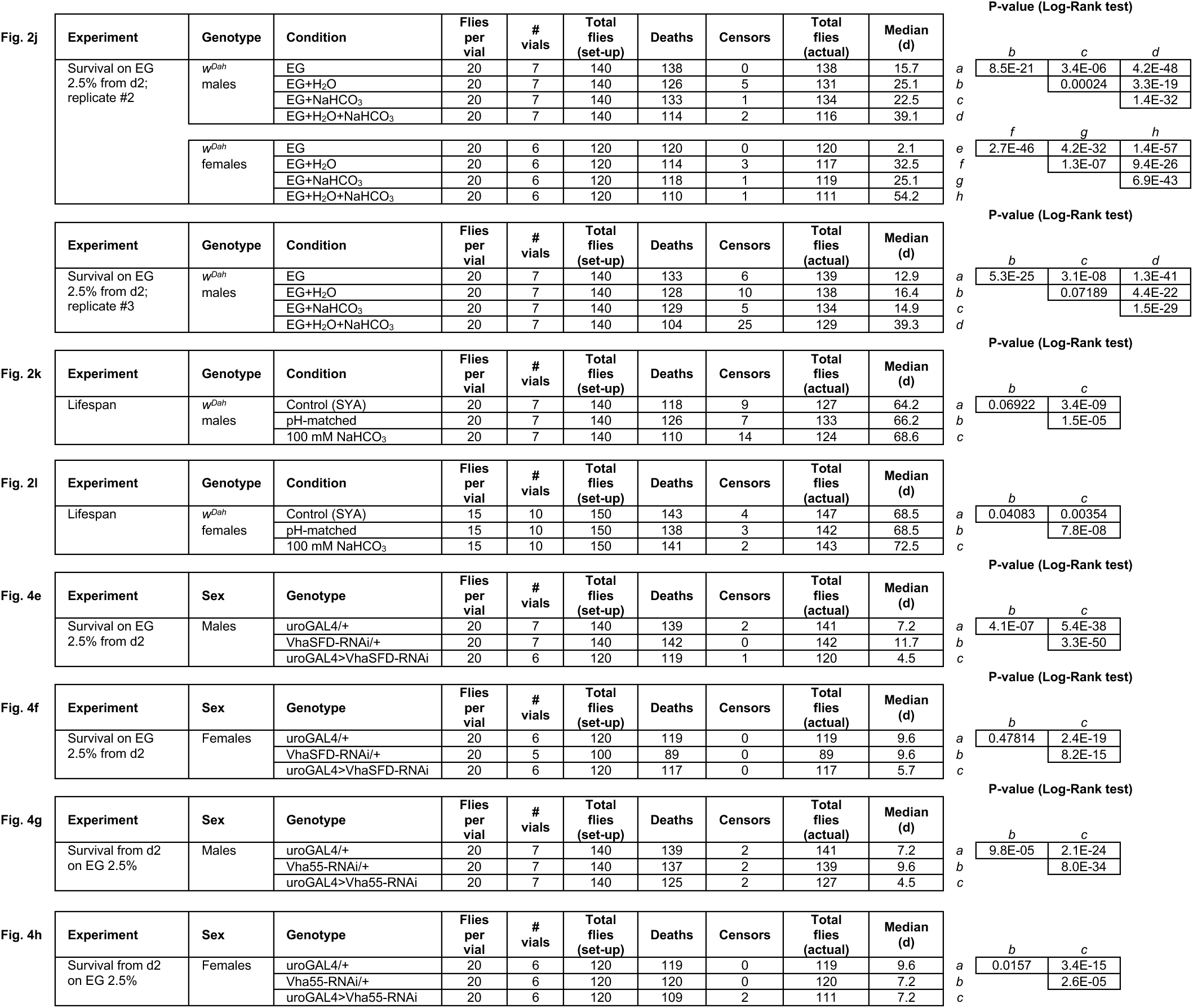
Summary of survival data. (Main manuscript figures)

**Supplementary Table 2.**
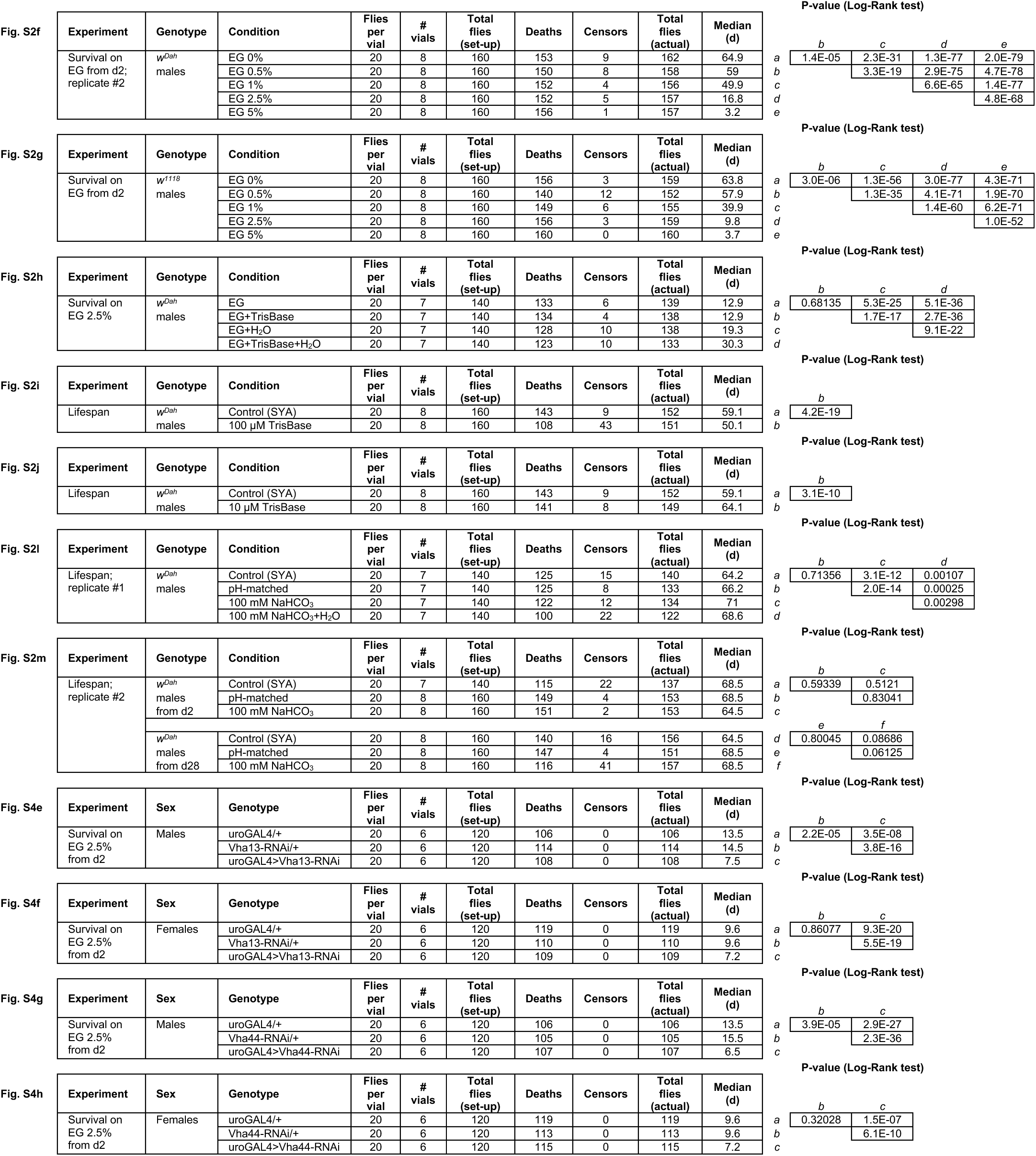
Summary of survival data. (Extended data figures)

